# Cis-regulatory elements driving motor neuron-restricted viral payload expression within the mammalian spinal cord

**DOI:** 10.1101/2024.09.02.610875

**Authors:** M. Aurel Nagy, Spencer Price, Kristina Wang, Stanley P. Gill, Erika Ren, Lorna McElrath, Victoria Pajak, Sarah Deighan, Bin Liu, Xiaodong Lu, Aissatou Diallo, Shih-Ching Lo, Robin Kleiman, Christopher Henderson, Junghae Suh, Eric C. Griffith, Michael E. Greenberg, Sinisa Hrvatin

## Abstract

Spinal motor neuron (MN) dysfunction is the cause of a number of clinically significant movement disorders. Despite the recent approval of gene therapeutics targeting these MN-related disorders, there are no viral delivery mechanisms that achieve MN-restricted transgene expression. In this study, chromatin accessibility profiling of genetically defined mouse MNs was used to identify candidate cis-regulatory elements (CREs) capable of driving MN-selective gene expression. Subsequent testing of these candidates identified two CREs that confer MN-selective gene expression in the spinal cord as well as reduced off-target expression in dorsal root ganglia. Within one of these candidate elements, we identified a compact core transcription factor (TF)-binding region that drives MN-selective gene expression. Finally, we demonstrate that selective spinal cord expression of this mouse CRE is preserved in non-human primates. These findings suggest that the generation of cell-type-selective viral reagents, in which cell-type-selective CREs drive restricted gene expression, will be valuable research tools in mice and other mammalian species, with potentially significant therapeutic value in humans.

**SIGNIFICANCE STATEMENT:** Motor neurons transduce the motor outputs of nervous system activity to muscle and are the vulnerable neurons in a number of clinically significant degenerative conditions, including spinal muscular atrophy and amyotrophic lateral sclerosis. A recent strategy for treating motor neuron degenerative diseases has been to use viruses to introduce genes into motor neurons to inhibit the degenerative process. However, we still lack viral reagents that promote gene expression in motor neurons without potentially toxic off-target expression in other cell types. Our study identifies cis-regulatory elements capable of conferring motor neuron-selective transgene expression in a viral context. These findings have important implications for future gene therapeutics for motor neuron-related disorders.

## INTRODUCTION

Motor neurons (MNs) constitute a major output of the mammalian central nervous system (CNS), controlling both voluntary and involuntary muscle movements (1). In addition, MN dysfunction underlies a number of severe, progressive human degenerative conditions, including spinal muscular atrophy (SMA) and amyotrophic lateral sclerosis (ALS) (2). Extensive investigation over decades in a variety of organisms has illuminated aspects of MN developmental specification and function. However, despite the recent regulatory approval of virus-based gene therapeutics targeting the progressive MN loss in SMA (3), to date, the ability to gain specific genetic access to MNs in mammals using engineered viral vector approaches has been limited (4–8).

In recent years, gene delivery via nonpathogenic replication-deficient recombinant adeno-associated virus (AAV) has been increasingly employed in research efforts due to a number of significant advantages, including utility in a broad range of mammalian species, transduction of diverse cell types, and easy experimental implementation. Moreover, the low immunogenicity of AAV relative to other common viral vectors and the minimally integrating nature of this virus has elevated this approach for clinical gene therapy applications. Indeed, five AAV-based treatments have been approved for commercialization to date in the United States, with dozens more candidates actively undergoing clinical trial (3, 9–12).

Despite these clinical advances, the vectors employed thus far largely utilize a combination of naturally derived viral serotypes and strong, ubiquitous promoters to maximize transgene expression in the target cell population. However, these initial constructs remain unoptimized with respect to minimizing transgene expression in off-target cell types. This shortcoming has manifested as adverse events in nonhuman primate (NHP) AAV preclinical studies, including dorsal root ganglion (DRG) mononuclear cell infiltration and sensory neuron degeneration following systemic or intrathecal AAV administration, presumably due to high off-target transgene expression in this cell type (13–16). Thus, the development of vectors driving more targeted transgene expression promises significant gains in therapeutic safety and efficacy.

While the ability of cis-acting gene regulatory elements to confer cell-type-specificity to viral transgene expression has long been recognized (17–21), recent advances in chromatin profiling techniques have enabled genome-wide comparative analyses across cell types to identify compact, gene-distal cis-regulatory elements (CREs) whose activity is restricted to particular cell types. Incorporation of these enhancer elements into recombinant AAVs, used in conjunction with viral capsid engineering methods, has begun to yield viral reagents with impressive specificity for the cell types of interest (22–27). However, despite the high clinical importance of lower MNs in the spinal cord and known liver- and DRG-derived toxicity observed in human MN-related disorder gene therapies, no CREs capable of restricting transgene expression to this cell population have been reported.

Here, we employ a genetically targeted chromatin profiling strategy to generate a genome-wide dataset of chromatin accessibility and gene expression in lower motor neurons of the spinal cord in the adult mouse. We select and functionally evaluate a subset of these CREs to identify two novel elements that drive MN-selective transgene expression in the spinal cord and achieve substantial reduction of transgene expression in DRG compared to the widely utilized CAG promoter. For one promising candidate CRE, we identify a compact core region capable of reproducing the selectivity of the full-length sequence with reduced packaging size and characterize key transcription factor (TF)-binding motifs that confer this motor neuron-selective activity. Finally, we demonstrate the preservation of spinal cord-selective payload expression of the top CRE and its compact core region in non-human primate. Together, our findings suggest that cell-type-selective viral reagents will be valuable research tools in mice and likely higher-order mammalian species, with potentially significant therapeutic value in humans.

## RESULTS

### Candidate cis-regulatory element identification and selection

To identify cis-regulatory elements (CREs) selectively active in motor neurons (MNs), spinal motor neuron nuclei were tagged and immunopurified using the INTACT (Isolation of Nuclei Tagged in specific Cell Types) system **(Figure 1a)** (28). To this end, *Choline acetyltransferase (Chat)-Cre* mice were crossed with the conditional *Sun1-sfGFP-6xMyc* mouse line, so as to stably mark the nuclear envelope of *Chat*-expressing cells with green fluorescent protein (GFP) (29, 30). In the spinal cord, the *Chat*-expressing cell population comprises skeletal motor neurons as well as visceral MNs and cholinergic interneuron populations (31–33). Confocal microscopy confirmed GFP expression in all MN subtypes of these *Chat-Cre; Sun1-sfGFP-6xMyc* (Sun1-ChAT) animals, including skeletal MNs, which were identified by their distinctive large somata, positive ChAT co-staining, and anatomic localization to the ventral horn **(Figure 1b).**

**Figure 1.**
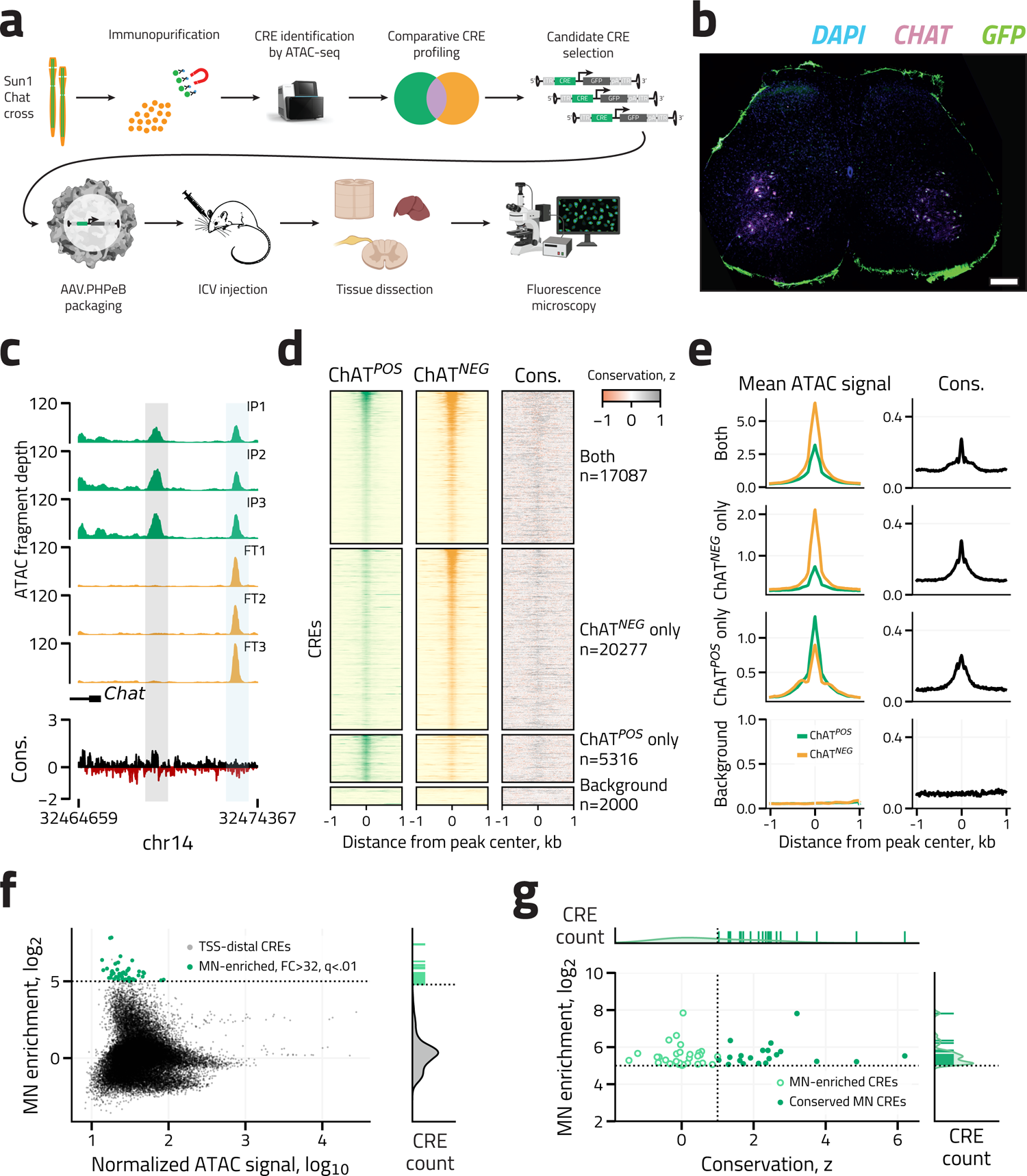
Experimental strategy and motor neuron CRE identification. **(a)** Schema of overall experimental design. **(b)** Representative IHC image of a L2 spinal cord section from a 10-week-old Sun1-ChAT mouse, demonstrating cytoplasmic ChAT (magenta) and GFP-labeled MN nuclear envelope (green). DAPI nuclei labeled in blue. Scale bar 250 μm. **(c)** Representative ATAC-seq genome browser tracks (normalized counts per location) across three bioreplicates (each bioreplicate comprises combined spinal cord tissue from two animals) derived from immunopurified spinal motor neuron (IP, green) and flow-through (FT, yellow) nuclei near the 5’ *Chat* gene terminus (blue). An example CRE that is selectively accessible in MNs (CRE98, gray highlight) and a nearby nonselective CRE (blue highlight) are shown. Sequence conservation across the Placental mammalian clade (PhyloP) is also shown. **(d)** Fixed-line-plots showing mean ATAC-seq signal strength (left and middle) and sequence conservation Z-score (PhyloP, placental mammals, right) in immunopurified spinal MN (green) and flow-through (yellow) nuclei, sorted by descending signal strength, across three bioreplicates (each bioreplicate comprises combined spinal cord tissue from two animals). Each ATAC-seq site is represented as a single horizontal line centered at the peak summit with flanking 1 kb. Color denotes ATAC-seq signal intensity (displayed in log_2_(counts per million + 1)). Sites are classified by differential accessibility across the immunoprecipitated spinal MN and flowthrough populations as follows: Both, accessible in both cell populations and not significantly different (FDR-corrected q > 0.05); ChAT*^NEG^*only, significant enrichment of accessibility in the flow-through population (ChAT*^POS^* / ChAT*^NEG^* fold-change ≤ 0.5, FDR-corrected q < 0.05, DESeq2); ChAT*^POS^* only, significant enrichment of accessibility in the spinal MN population (ChAT*^POS^*/ ChAT*^NEG^* fold-change ≥ 2, FDR-corrected q < 0.05, DESeq2); Background (n = 2,000), randomly selected genomic loci included for visual comparison. **(e)** Aggregate ATAC signal (left) and conservation (right) across peaks described in Figure 1d plotted as mean ATAC signal by distance from peak center. **(f)** Left, MA plot of MN-enrichment (log2(ChAT*^POS^*/ChAT*^NEG^* ATAC-seq signal) as a function of mean ATAC-seq signal for each TSS-distal peak. Sites that exhibit >32-fold enriched accessibility in MNs with FDR-corrected q < 0.01 are denoted in green. Right, histogram of MN enrichment for all peaks with MN-enriched peaks denoted by green rug plot (>32-fold enriched, q < 0.05). **(g)** Left, scatter plot of MN-enrichment (log2(ChAT*^POS^*/ChAT*^NEG^* ATAC-seq signal) as a function of conservation Z-score (PhyloP, placental mammals) for each MN-enriched (>32-fold enriched accessibility in MNs, FDR-corrected q < 0.01) TSS-distal peak (light green hollow points). Sites with conservation Z-score > 1 denoted in dark green filled points. Top and Right, corresponding histograms across conservation z-score and MN-enrichment, respectively.

Cell identity was also verified transcriptionally following immunopurification of tagged nuclei. Bulk RNA-seq of immunopurified (ChAT*^POS^*) as well as putatively MN-depleted flowthrough (ChAT*^NEG^*) nuclei was performed to identify genes differentially expressed between these two populations. As expected, the cholinergic marker genes *Slc5a8* and *Chat* were enriched in the ChAT*^POS^* population relative to ChAT*^NEG^* nuclei. By contrast, expression was diminished for glutamatergic (*Slc17a8*, *Slc17a6*) and GABAergic (*Gad1*, *Slc6a5*) neuron, oligodendrocyte (*Mbp*, *Mobp*), astrocyte (*Gfap*, *Aqp4*), microglia (*Cx3cr1*, *Trem2*), and endothelial cell (*Cldn5*) marker genes (30–32, 34), confirming successful purification of *Chat*-expressing nuclei relative to the other major spinal cord cell types **(Supplementary Figure 1a)**. To further distinguish between *Chat*-expressing subpopulations, the relative enrichment of skeletal MN *(Bcl6*, *Ahnak2*, *Aox1)*, visceral MN (*Mme*, *Gnb4*, *Nos1*), and cholinergic interneuron (*Pou6f2*, *Pax2*, *Ebf2*) marker genes was assessed across the ChAT*^POS^* and ChAT*^NEG^*purified nuclear populations (30). Only skeletal MN markers demonstrated significant enrichment in the ChAT*^POS^* population (q-value < .01, FC > 2, DESeq2) over ChAT*^NEG^*samples, further confirming that skeletal MNs comprised the majority of purified nuclei **(Supplementary Figure 1b, Materials and Methods),** consistent with previous studies employing this approach (30). These data provide additional genetic characterization of adult murine motor neurons and are made available as a resource with this publication (GSE accession to be determined).

Enhancers are gene-distal DNA regulatory elements that confer cell-type- and context-specific expression through selective transcription factor recruitment (35, 36). The resulting local nucleosome depletion enables candidate CRE identification with genome-wide nucleosome profiling techniques. Having verified the predominantly skeletal MN identity of our ChAT*^POS^*population, ATAC-seq (assay for transposase accessible chromatin with sequencing) (37) was employed to identify nucleosome-depleted CREs in ChAT*^POS^* and ChAT*^NEG^*nuclei (n = 22,403 and 37,365 peaks, respectively; union n = 42,680 peaks, representing all identified putative CREs in these populations) **(Figure 1c-e, Materials and Methods)**. Standardized quality control metrics, including nucleosomal ATAC-seq fragment size distribution, high irreproducible discovery rate (IDR), and sample correlation were assessed to confirm the technical soundness and internal consistency of ATAC-seq data from this sparse cell population **(Supplementary Figure 1c-e)** (38).

Since the relative chromatin accessibility of a CRE predicts its potential activity as a functional regulator of gene expression (39), putative CREs were ranked by their selective local chromatin accessibility in the ChAT*^POS^* population relative to ChAT*^NEG^* (40). As enhancers are gene distal, only peaks >250 base pairs (bp) from transcriptional start sites (TSSs) with at least 32-fold enrichment for ChAT*^POS^* nuclei relative to ChAT*^NEG^* were selected, yielding 49 putative CREs for further evaluation (FDR-corrected q < 0.05) **(Figure 1f)**. To enrich for CREs that could be useful across mammalian species, the most evolutionarily conserved elements (conservation z-score across the placental clade > 1, **Methods**) were sub-selected from this population to obtain a set of 20 high-likelihood, MN-restricted enhancers for downstream functional evaluation (**Figure 1g**).

### AAV fluorescent reporter imaging identifies motor neuron-enriched CREs

Given limitations associated with the sparsity of skeletal MNs, we eschewed a large-scale screening approach and employed traditional techniques to test the most promising CRE candidates individually for their ability to drive MN-restricted expression of a GFP transgene in an AAV reporter construct. To this end, from among the 20 high-confidence MN-restricted CREs, we selected elements exhibiting the largest fold-enrichment of MN accessibility (CRE187, CRE219, CRE150), the greatest statistical significance for MN-enriched accessibility (CRE98, CRE32, CRE226), and the highest degree of mammalian conservation (CRE226, CRE057, CRE119) for evaluation. As negative controls, we also included three CREs of similar genomic size that performed uniformly poorly across these metrics (CRE58, CRE70, CRE76), yielding a total of 11 elements for testing (CRE226 appeared in two categories, **Figures 2a, 2b)**.

**Figure 2.**
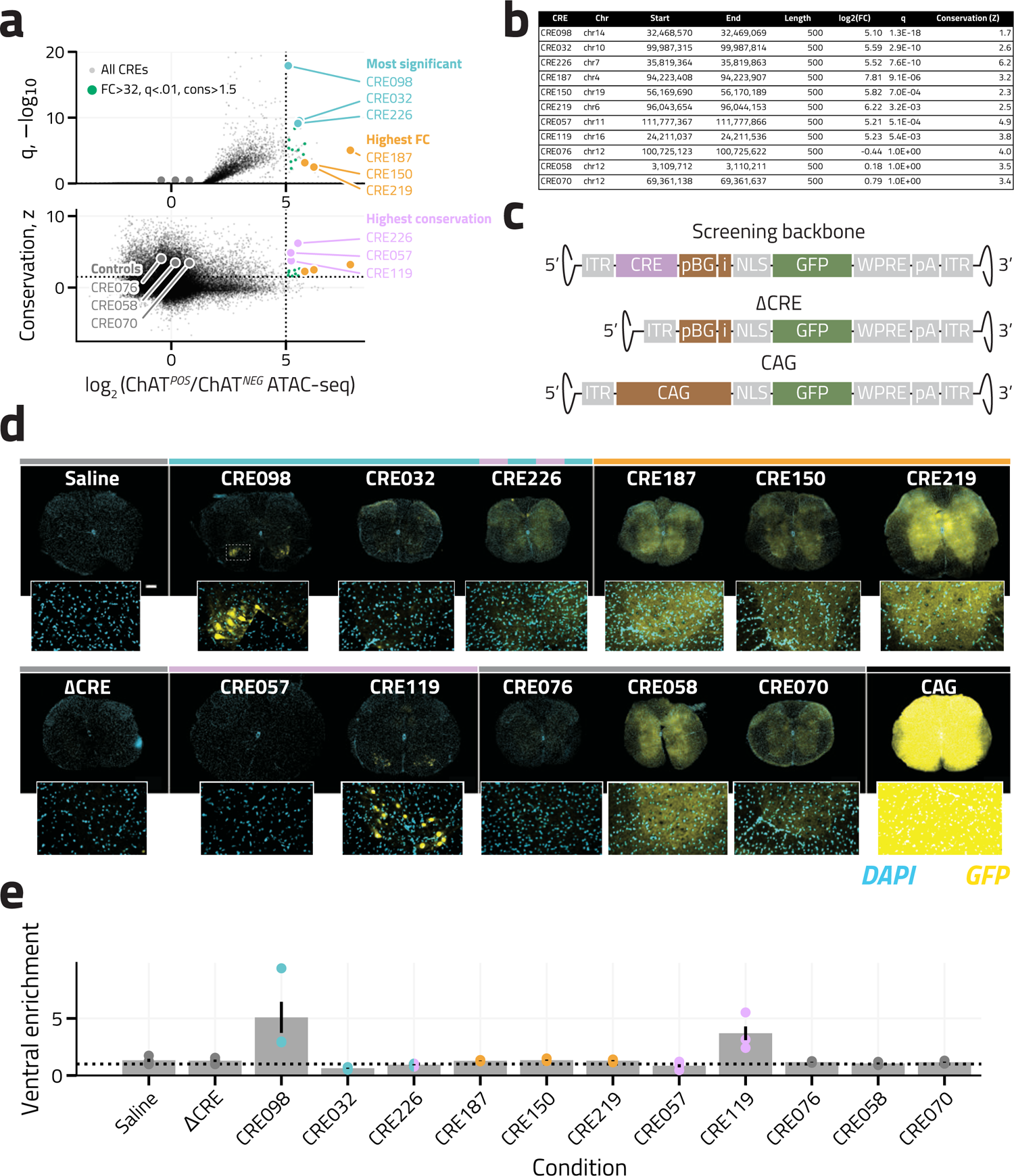
Initial candidate CRE screening by confocal microscopy. **(a)** Upper panel: Volcano plot of –log_10_(FDR-corrected q) versus MN-enrichment (fold change, log2[ChAT*^POS^*/ChAT*^NEG^* ATAC-seq signal]) for all TSS-distal putative CREs. Lower panel: Conservation z-score versus MN-enrichment (fold change, log2[ChAT*^POS^*/ChAT*^NEG^* ATAC-seq signal]) for all TSS-distal putative CREs. Vertical dotted line denotes 32-fold MN enrichment threshold, horizontal dotted line denotes conservation Z-score threshold of 1. Putative MN-enriched (>32-fold enriched accessibility in MNs, FDR-corrected q < 0.01) and conserved (Z-score > 1) CREs plotted in green. CREs selected from this subpopulation for final screening are highlighted and labeled by their relevant selection criteria: lowest q-value in blue, highest MN enrichment in yellow, most conserved in purple, and controls in gray. **(b)** Table of candidate CREs selected for testing, showing internal identification (CRE), genomic position (Chromosome, Start, End), element length (defined as 500 by peak-calling algorithm), MN ATAC-seq enrichment (calculated as log2(ChAT*^POS^*/ChAT*^NEG^* ATAC-seq signal, labeled as log2[FC]) and associated FDR-corrected q-value (q), and conservation Z-score. **(c)** AAV screening construct viral genome maps (not drawn to scale). ITR, inverted terminal repeats; pBG, minimal beta-globin promoter; i, intron; NLS, nuclear localization sequence; GFP, green fluorescent protein. WPRE, woodchuck hepatitis virus post-transcriptional response element; pA, poly-A tail. **(d)** Representative confocal native fluorescence images of T1-L4 spinal cord sections from wild-type mice 14 days after P0 ICV injection of candidate CRE reporter AAVs (DAPI, cyan; GFP, yellow). CRE selection criteria denoted by line above image labels (lowest MN differential ATAC-seq q-value in blue, highest MN ATAC-seq enrichment in yellow, most conserved in purple, controls in gray, and CAG in black). Scale bar 250 μm. Combined ventral horn images also shown at higher magnification in inset (denote by dashed box, 4.7x magnification). **(e)** Quantification of GFP signal intensity fold-change in ventral compared to dorsal horn for all constructs tested (ventral enrichment). Mean enrichment across three spinal cord sections per animal denoted by points, mean across three animals denoted by bar, standard deviation across three animals denoted by line. CRE selection criteria denoted by point color (lowest MN differential ATAC-seq q-value in blue, highest MN ATAC-seq enrichment in yellow, most conserved in purple, controls in gray, and CAG in black). Null hypothesis (ventral enrichment = 1) plotted as dotted line.

CREs were amplified from C57BL/6 mouse genomic DNA and incorporated into a nuclear localized GFP reporter (nlsGFP) AAV2 vector backbone (5’-ITR-CRE-pBG-nlsGFP-WPRE-polyA-ITR-3’) driven by a minimal *beta-globin* promoter (pBG), as described previously (22) **(Figure 2c)**. In parallel, we generated a negative control pBG construct lacking a CRE (ΔCRE) as well as a positive control construct in which GFP expression was driven by the strong, ubiquitously active CAG promoter (9), so as to define lower and upper bounds of payload expression. The resulting constructs were then individually packaged into the PHP.eB AAV capsid, which efficiently transduces spinal MNs after intracerebroventricular (ICV) administration in mouse (41, 42).

Wild-type C57BL/6 mice (n = 2-3 per condition) were singly dosed with 1.2 x 10^11^ viral genome copies (vgc, 4 μL) of AAV at postnatal day 0 (P0) via unilateral ICV injection. Saline injections were also performed as a control to establish baseline autofluorescence (n=3). Two weeks post-injection, thoracic and lumbar spinal cords (T4-L4) were dissected, transversely sectioned, and native DAPI and GFP expression (n = 2-5 sections per animal) were assessed by fluorescence microscopy. Relative skeletal MN enrichment was calculated by obtaining the ratio of GFP fluorescence intensity overlying DAPI-stained nuclei in the anatomically defined ventral and dorsal horns, where skeletal lower motor and second-order sensory spinal neuron somata are found, respectively. Notably, two of the eleven CRE-driven constructs drove increased ventral over dorsal horn GFP expression (CRE98: 5.09 ± 1.36-fold, CRE119: 3.70 ± 0.59-fold) compared to saline (1.28 ± 0.10-fold) and ΔCRE (1.33 ± 0.14-fold) (**Figures 2d, 2e, Supplementary** Figures 2a-c). CAG drove significant on- and off-target expression in the spinal cord, confirming the known poor skeletal MN-specificity of this promoter.

### Quantitative CRE evaluation by immunohistochemistry of AAV fluorescent reporter expression

To more precisely evaluate the efficiency and uniformity of CRE-driven gene expression relative to CAG, GFP expression was quantified via immunohistochemistry (IHC) across on-target ventral MNs (hereafter defined by positive co-staining of ChAT and the neuronal marker NeuN) and off-target other spinal neurons (defined by NeuN-positive staining without ChAT) for the following subset of constructs: saline (technical control), CRE98 and CRE119 (top hits), CRE57 and ΔCRE (negative controls), and CAG. Mice were injected and dissected as previously described for IHC and confocal imaging (**Methods**). Representative axial spinal cord sections for each construct after IHC-based labeling of NeuN, ChAT, and GFP are shown in **Figure 3a**. Two quantitative measures were evaluated for each construct: expression strength (mean GFP signal intensity overlying DAPI-stained nuclei for each cell in a given slice, **Supplemental Figures 3a**) and transduction efficiency (the percentage of transduced neurons with each neuron classified as GFP*^POS^* if cell signal intensity was greater than threshold, defined as the 99^th^ percentile of GFP signal intensity in neurons of the control saline condition, **Supplemental Figure 3b**, **Methods**). As expected, there was no significant difference in GFP expression across on- and off-target spinal neuronal populations for any of the CRE57-, ΔCRE-, and saline-injected control conditions by either of these metrics (**Supplemental Figure 3c, 3d)**.

**Figure 3.**
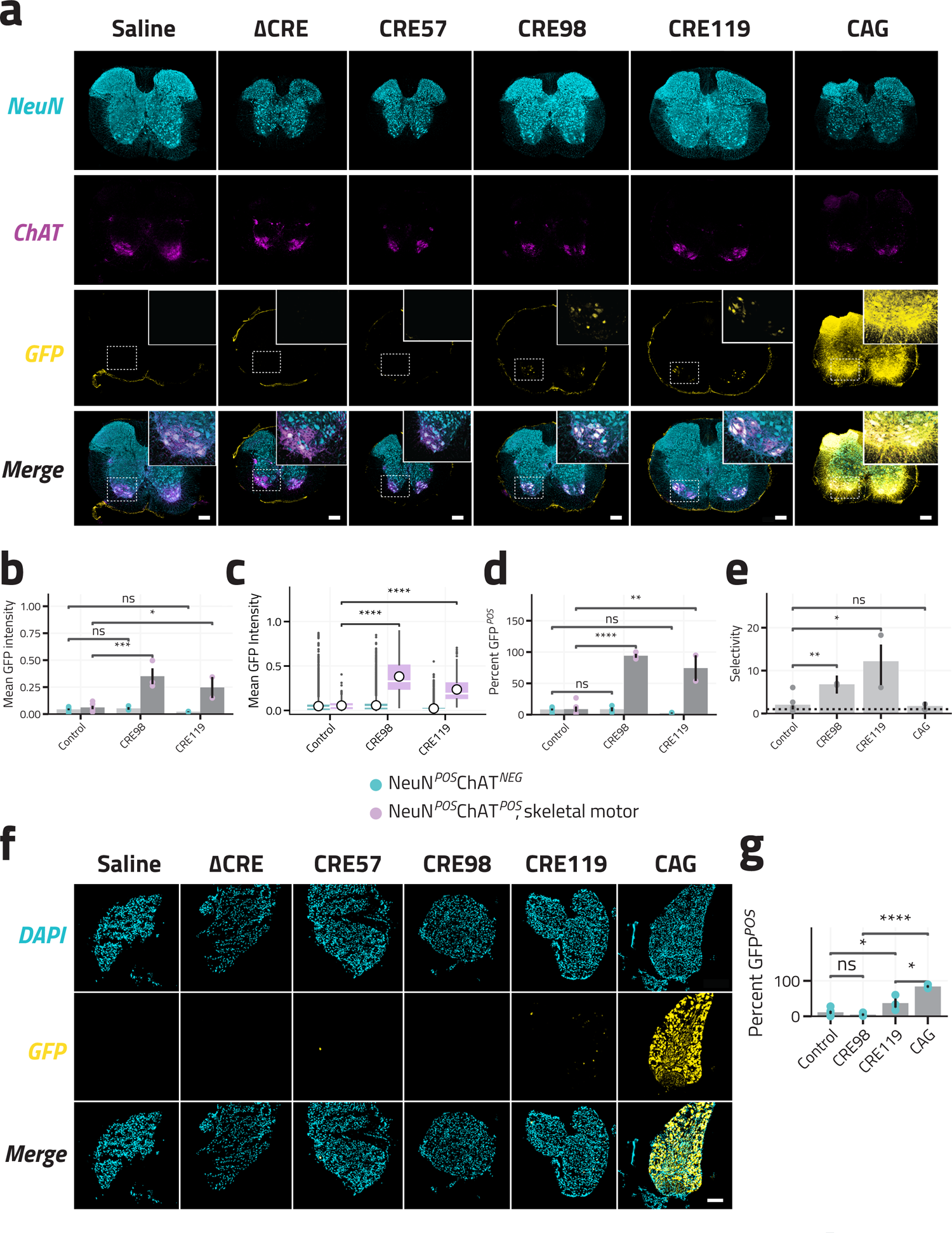
Quantification of CRE specificity by immunohistochemistry. **(a)** Representative IHC images of T1-L4 spinal cord sections 14 days after P0 ICV injection of candidate CRE reporter AAVs (NeuN, cyan; ChAT, magenta; GFP, yellow. Scale bar 250 μm. GFP and Merge images of ventral horn also shown as magnified inset). **(b)** Quantification of mean GFP signal intensity in off-target spinal neurons (NeuN*^POS^* ChAT*^NEG^*, blue) and on-target spinal motor neurons (NeuN*^POS^* ChAT*^NEG^*, purple) for combined CRE57/ΔCRE control (n = 6 mice), CRE98 (n = 3 mice), and CRE119 (n = 2 mice) across animals and **(c)** across all cells. Mean enrichment across all spinal cord sections per animal denoted by each point, mean across all animals per condition denoted by bar plot, standard deviation across all animals denoted by line. **(d)** Percentage of GFP*^POS^* cells in off-target spinal neurons (NeuN*^POS^* ChAT*^NEG^*, blue) and on-target spinal motor neurons (NeuN*^POS^*ChAT*^POS^*, purple) for combined CRE57/ΔCRE control (n = 6 animals), CRE98 (n = 3 animals), and CRE119 (n = 2 animals) constructs. Mean percentage across all spinal cord sections per animal denoted by each point, mean across all animals per condition denoted by bar plot, standard deviation across three animals denoted by line. **(e)** Selectivity (on-target NeuN*^POS^*ChAT*^POS^* / off-target NeuN*^POS^* ChAT*^NEG^*mean signal intensity across all neurons per animal) for combined CRE57/ΔCRE control (n = 6 animals), CRE98 (n = 3 animals), CRE119 (n = 2 animals), and CAG (n = 3 animals) constructs. Mean percentage across all spinal cord sections per animal denoted by each point, mean across all animals per condition denoted by bar plot, standard deviation across all animals denoted by line. **(e)** Quantification of GFP signal intensity in on-target spinal motor neurons (NeuN*^POS^* ChAT*^NEG^*) for combined saline/ΔCRE/CRE57 controls, CRE098, and CRE119 across all cells (combined across all animals). Mean intensity across all cells per condition denoted by white circle, distribution of all cells by section denoted by box-and-whisker plot (median denoted by white line, first and fourth quartiles by vertical line, outliers by black points). Images acquired and quantified at pBG promoter-optimized acquisition parameters. **(f)** Representative IHC images of T1-L4 DRG simultaneously dissected from previously described animals 14 days after P0 ICV injection of candidate CRE reporter AAVs (DAPI, cyan; GFP, yellow). Scale bar 100 μm. **(g)** Percentage of GFP*^POS^* off-target DRG neurons (visually identified by nuclear DAPI morphology and size) for combined CRE57/ΔCRE control (n=6), CRE98 (n=3), CRE119 (n=2), and CAG (n=3) constructs. Mean percentage across all DRG sections per animal denoted by each point, mean across all animals per condition denoted by bar plot, standard deviation across all animals denoted by line. All images acquired at pBG promoter-optimized acquisition parameters. Significance measured by Bonferroni-corrected two-tailed t test. *** q < 0.001, ** q < 0.01, * q < 0.05, *ns* q > 0.05.

Mean on-target GFP expression strengths were significantly greater for CRE98 and CRE119 compared to aggregated saline/CRE57/ΔCRE controls when comparing across animals (5.7-fold, p = 9.5e-4, and 4.0-fold, p = 1.4e-2 respectively; T-test, BH-corrected, **Figure 3b**) or all cells (7.0-fold, p=2.6e-182, and 4.3-fold, p = 9.9e-96 respectively; T-test, BH-corrected, **Figure 3c**). Both CRE98 and CRE119 also drove significantly more efficient on-target expression (94.0 ± 3.2% GFP*^POS^*, p = 3.4e-6 and 74.3 ± 20.6% GFP*^POS^*, p = 1.8e-3 respectively) compared to aggregated controls (8.2 ± 1.6% GFP*^POS^*; T-test, BH-corrected, **Figure 3d**). By contrast, off-target spinal neuron CRE98- and CRE119-driven reporter expression was not significantly different from controls when assessed by either signal intensity (1.2-fold, p = 5.8e-1 and 0.5-fold, p = 2.0e-1, respectively; T-test, BH-corrected; **Figure 3b**) or transduction efficiency (8.4 ± 3.2%, p = 9.4e-1 and 2.4 ± 0.9%, p = 9.1e-1 respectively) compared to the control condition (8.2 ± 1.6%; T-test, BH-corrected; **Figure 3d**). Together, these findings demonstrate that CRE98- and CRE119-driven constructs achieve selective and efficient on-target expression in the spinal cord MNs.

We also quantified these metrics for the CAG-driven construct (representative axial spinal cord sections after IHC acquired at both pBG- and CAG-optimized intensity windowing parameters shown in **Supplementary Figure 3e** and quantified in **Supplementary Figure 3f, Methods)**. Relative to the ΔCRE control condition, on-target MN expression was 12.0-fold increased (p = 9.8e-2; T-test, BH-corrected, **Supplementary Figure 3g**), with 100% transduction efficiency (**Supplementary Figure 3h**; threshold defined by 99%-tile of ΔCRE expression, **Supplementary** Figures 3i), confirming the known high expression of CAG-driven vectors in motor neurons of the spinal cord. However, off-target spinal neuron expression was also 7.1-fold greater (p = 3.2e-2, T-test, BH-corrected) than that observed in the control condition (**Supplementary Figure 3g**), and 100% of off-target neurons were also classified as GFP*^POS^* by the previously described approach (**Supplementary Figure 3h**). These findings confirm that relative to the candidate CRE-driven constructs, the increased on-target CAG-driven expression comes with substantial off-target spinal neuron expression.

Finally, we examined the ratio of mean on- to off-target signal intensities for each construct as a proxy for selectivity (**Figure 3e**). CRE98- and CRE119-driven selectivity were significantly increased (11.4-fold, p = 2.1e-2 and 6.7-fold, p = 7.0e-3, respectively; T-test, BH-corrected) compared to aggregate controls (1.4-fold). By comparison, CAG demonstrated only 1.8-fold enhancement in motor neurons relative to other spinal neurons, not statistically different from controls (p = 8.4e-1, T-test, BH-corrected). Together, these findings highlight the improved specificity of these CRE constructs for motor neurons of the spinal cord over the standard CAG promoter.

### CRE-driven viral transgene expression outside the spinal cord

In clinical contexts, off-target AAV payload expression can introduce safety concerns that limit viral vector doses and impede therapeutic efficacy. Such toxicity is dependent on the route of administration, but clinically notable examples include dorsal root ganglion (DRG) and hepatotoxicity via intrathecal and intravenous administration, respectively (43, 44). We therefore examined the extent to which CRE98 and CRE119 restrict off-target expression in mouse thoracic/lumbar DRGs simultaneously dissected from the ICV-injected animals described in the prior section. As before, virus-derived GFP expression rates in these off-target tissues were quantified by classifying cells as GFP*^POS^* if IHC signal intensity overlying DAPI-stained nuclei was greater than the 99th percentile of cells in the saline condition (**Methods**).

Significantly fewer DRG neurons exhibited GFP expression in animals injected with the CRE98 construct (4.5 ± 3.1%) compared to CAG (84.1 ± 3.2%, 0.05-fold, p = 5.4e-5; T-test, BH-corrected; **Figures 3f, 3g)**, and CRE98-driven GFP expression did not differ from controls in this off-target tissue (p = 3.3e-1, T-test, BH-corrected). CRE119 drove an intermediate level of expression in DRG neurons (37.0 ± 12.5%, 3.32-fold, p = 3.9e-2; T-test, BH-corrected) relative to controls, though this was still significantly decreased relative to CAG (0.44-fold, p = 2.2e-6; T-test, BH-corrected). Thus, at least with respect to the DRG, MN-selective CREs possess significantly reduced off-target tissue activity relative to the widely employed CAG promoter.

### Native genomic contexts of CRE98 and CRE119

Having identified two mammalian CREs that impart skeletal MN-selective expression when applied in the AAV context in mouse, we next examined the native genomic context of these CREs to identify potential target genes. CRE98 is located approximately 3 kilobases (kb) upstream of two overlapping genes essential for cholinergic signal transduction: *Chat* and *Slc18a3 (*entirely contained within the first intron of *Chat*), respectively encoding the key enzyme for the synthesis of the motor neurotransmitter acetylcholine and its vesicular transporter (**Figure 1c**) (45). A homologous sequence to CRE98 has been previously characterized in rat, where *in vitro* analyses identified an upstream 2.3 kb element with both off-target silencing function (attributed to the restrictive element-1 silencing transcription factor [REST]) and on-target activating capabilities in transgenic cell lines (46, 47). To our knowledge, no prior work has demonstrated preservation of CRE98 function in viral heterologous contexts, or investigated whether this element is preserved or retains its function in larger vertebrates.

By contrast, there are no spinal MN-restricted genes or MN development- or function-associated genes within 800 kb of CRE119 (identified genes within this window include *Bcl6* [tumor suppressor], *Rtp2*/*Rtp4* [cell surface protein receptors], *Masp1* [leptin complement component), *Sst* [somatostatin neurotransmitter], and *Lpp* [gene of unknown function frequently fused in benign lipomas]). As this MN-selective CRE would not have been identified by testing loci proximal to known MN marker genes, our study highlights the importance of comparative chromatin accessibility profiling for unbiased CRE selection.

### Mechanistic investigation of mutant and synthetic CRE98 variants

To gain further insight into the mechanisms underlying MN-restricted CRE98 activity, we next sought to identify the sequence features contributing to its MN-selective expression. We generated a series of nuclear GFP reporter AAV constructs harboring truncated CRE98 sequence variants (**Figure 4a**). These were injected in mice, and the percentage GFP*^POS^* and GFP signal intensity were measured by IHC in on- and off-target spinal neurons as described above (**Figures 4b and 4c**, respectively). As observed previously, the full-length CRE98 construct (696 bp) drove high rates of GFP expression in the on-target spinal MN population (82.9 ± 6.0%), which was largely preserved in 5’ and 3’ deletion constructs (A: 70.9 ± 2.1%, p = 2.3e-1 and C: 66.2 ± 3.7%, p = 1.4e-1, respectively; T-test, BH-corrected), but not in further truncations lacking a central core region (B: 5.6 ± 5.6%, p = 3.1e-3, and D: 6.2 ± 3.1%, p = 3.4e-4, respectively; T-test, BH-corrected). Indeed, CRE98 variants containing only this central core region preserved MN-selective reporter expression (E: 76.6 ± 0.8%, p = 4.5e-1 and F: 78.2 ± 0.7%, p = 5.9e-1, respectively; T-test, BH-corrected) and signal intensity (9.5-fold and 10.3-fold enrichment in on-target skeletal MNs compared to off-target spinal neurons, respectively, p = 9.7e-1), indicating that this region is both necessary and sufficient for MN-selective expression.

**Figure 4.**
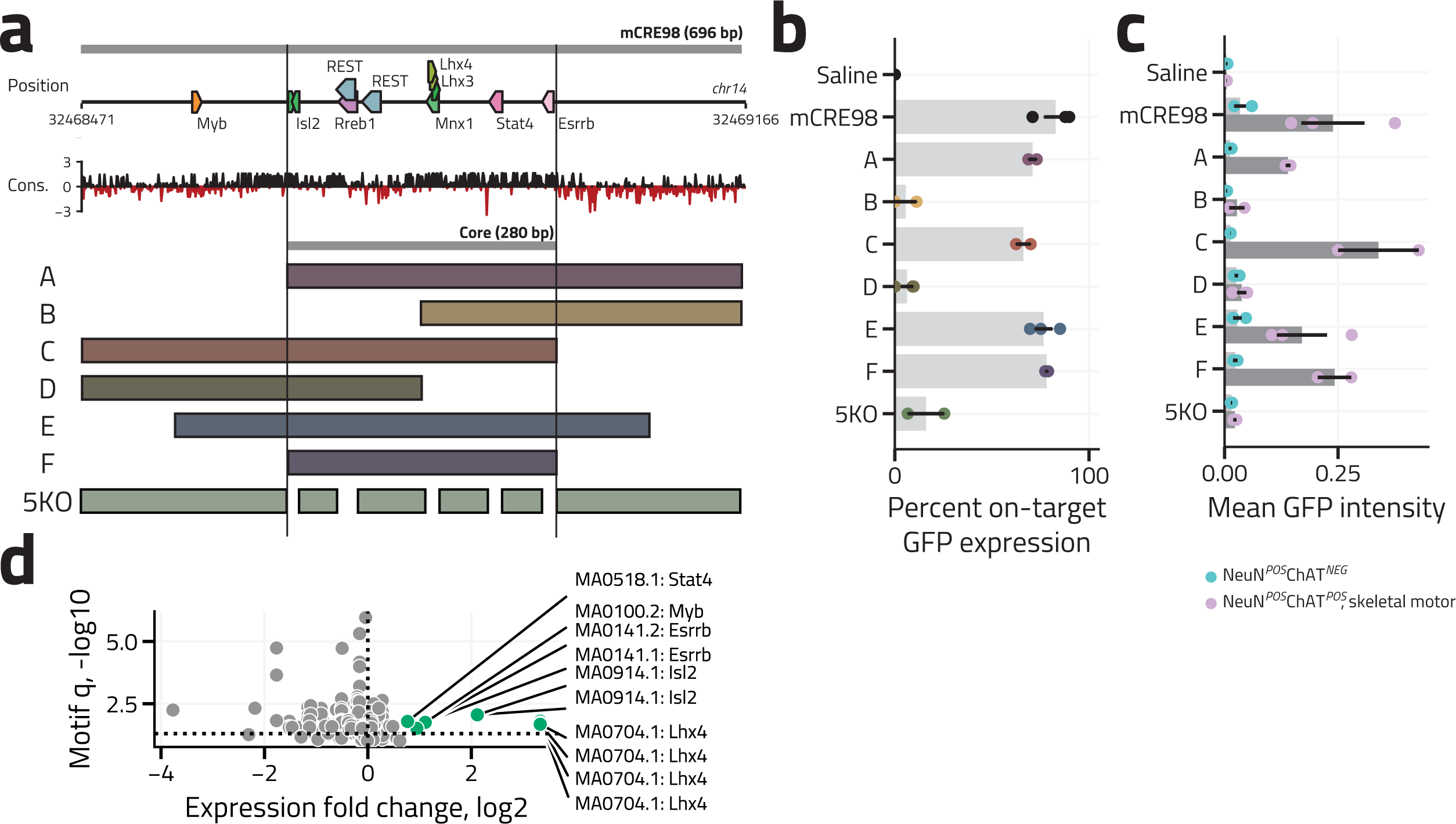
A core TF-binding region mediates CRE98 function. **(a)** Top: Native genomic context of mouse CRE98 (mCRE98) showing TF-binding sites (TFBS) along genomic coordinates of chromosome 14. Middle: Raw conservation score by base-pair conservation (PhyloP, placental clade). Bottom: CRE98 core region (mCORE), truncation constructs (A-F), and 5KO targeted deletion construct. **(b)** Percentage GFP*^POS^* cholinergic cells (ChAT*^POS^*) and **(c)** mean GFP signal intensity in off-target spinal neurons (NeuN*^POS^*ChAT*^NEG^*, blue) and on-target spinal motor neurons (NeuN*^POS^*ChAT*^POS^*, purple) for saline-injected control (n = 2 animals), murine CRE98 (n = 3 animals), and truncation and 5KO constructs (n = 2-3 animals per construct). Mean across all spinal cord sections per animal denoted by each point, mean across all animals per condition denoted by bar plot, standard deviation across three animals denoted by line. Images acquired and quantified at pBG promoter-optimized acquisition parameters. **(d)** Scatter plot of statistical significance (-log10 [Benjamini-Hochberg corrected TF motif enrichment q-value in CRE98)) as a function of MN-enrichment (log2 [ChAT*^POS^*/ChAT*^NEG^*RNA-seq signal]). Motif enrichment q-values derived from FIMO. Motifs that exhibit >2-fold expression enrichment in MNs with FDR-corrected q < 0.01, and motif-enrichment q < 0.01 are denoted in green and labeled by associated transcription factor.

CREs achieve cell-type-specific control of gene expression through the combinatorial recruitment of transcription factors (TFs) (48). We therefore interrogated the CRE98 sequence for known TF-binding motifs using the FIMO MEME analysis tool (**Methods**) (49), leveraging our skeletal MN RNA-seq data to identify TF-binding motifs whose corresponding TFs were selectively expressed in the purified ChAT*^POS^*nuclei (FDR-corrected q < 0.05) relative to the off-target ChAT*^NEG^* population (**Figure 4d**). This analysis identified Lhx4/Mnx1/Lhx3, Isl2, Esrrb, Myb, and Stat4 as candidate CRE98-regulating TFs **(Supplementary Figure 4a)**. These TFs, and/or related family members with similar binding motifs, have been demonstrated to either be MN-defining during differentiation or to serve as MN subtype markers in adult mice (30, 31). Two additional motifs were identified as high-likelihood functional sequences in CRE98 though they failed to meet the *a priori* MN-selective RNA-seq expression threshold: the Rreb1 motif (FDR-corrected q = 0.078), which has been implicated as a MN subclass-specific gene (32), and the REST motif, which was previously identified as necessary for restriction of reporter expression to neuronal cell lines by the homologous rat sequence (47). Notably, all identified motifs except for that of Myb were located within the central core region.

To investigate the contribution of these TF-binding sites, we generated a CRE98 mutant construct (5KO) harboring precise deletions of five of the six TF-binding sites identified in the above core region (Isl2, Lhx4/Mnx1/Lhx3, Stat4, Esrrb, and 1 of 2 REST motifs + Rreb1, **Figure 4a**). Compared to full-length CRE98, the 5KO construct demonstrated a profound decrease in MN-selective reporter gene expression (16.0 ± 9.4% of GFP*^POS^* ChAT*^POS^* cells, p = 7.7e-3, T-test, BH-corrected, **Figures 4b, 4c**). Notably, the loss of these binding sites in CRE98 did not promote aberrant payload expression in off-target spinal populations (**Figure 4c**), suggesting that the corresponding TFs likely confer MN selectivity via an activating mechanism in the appropriate cellular context. In summary, these findings suggest that disruption of these core Isl2-, Lhx4/Mnx1/Lhx3-, Stat3-, Esrrb-, and Rreb1-binding sites is sufficient to eliminate MN-selective CRE98 function, though it remains to be determined whether additional TFs also contribute to this effect.

Having identified a central core sequence that largely recapitulated full-length CRE98 function and key TF-binding sites necessary for its function, we next tested a reporter construct incorporating triplicate concatemers of the CRE98 core sequence (3mCORE), as well as a similar triplicate concatemer of a synthetic 129 bp sequence comprising just the core TF motifs separated by short 15 bp linkers (3mTFBS, **Figures 5a, 5c**). Relative to CRE98 (mean intensity 0.01 ± 0.0, 4.5 ± 3.1% GFP*^POS^*), the concatemerized CRE98 core construct increased reporter expression in both on-target skeletal MNs (mean intensity 0.80 ± 0.03 [**Figure 5d**], 100.0 ± 0.0% GFP*^POS^* [**Figure 5e**]) and off-target spinal (mean intensity 0.34 ± 0.07, 76.8 ± 14.7% GFP*^POS^*[**Supplementary Figures 5a, 5b**]) and DRG neurons (mean intensity 0.13 ± 0.05, 67.5 ± 11.2% GFP*^POS^*, **Supplemental Figure 5c-e**). By contrast, the fully synthetic 3mTFBS construct drove weak non-selective expression in spinal cord and DRG neurons, suggesting native motif positioning and spacing play an important role in conferring MN specificity.

**Figure 5.**
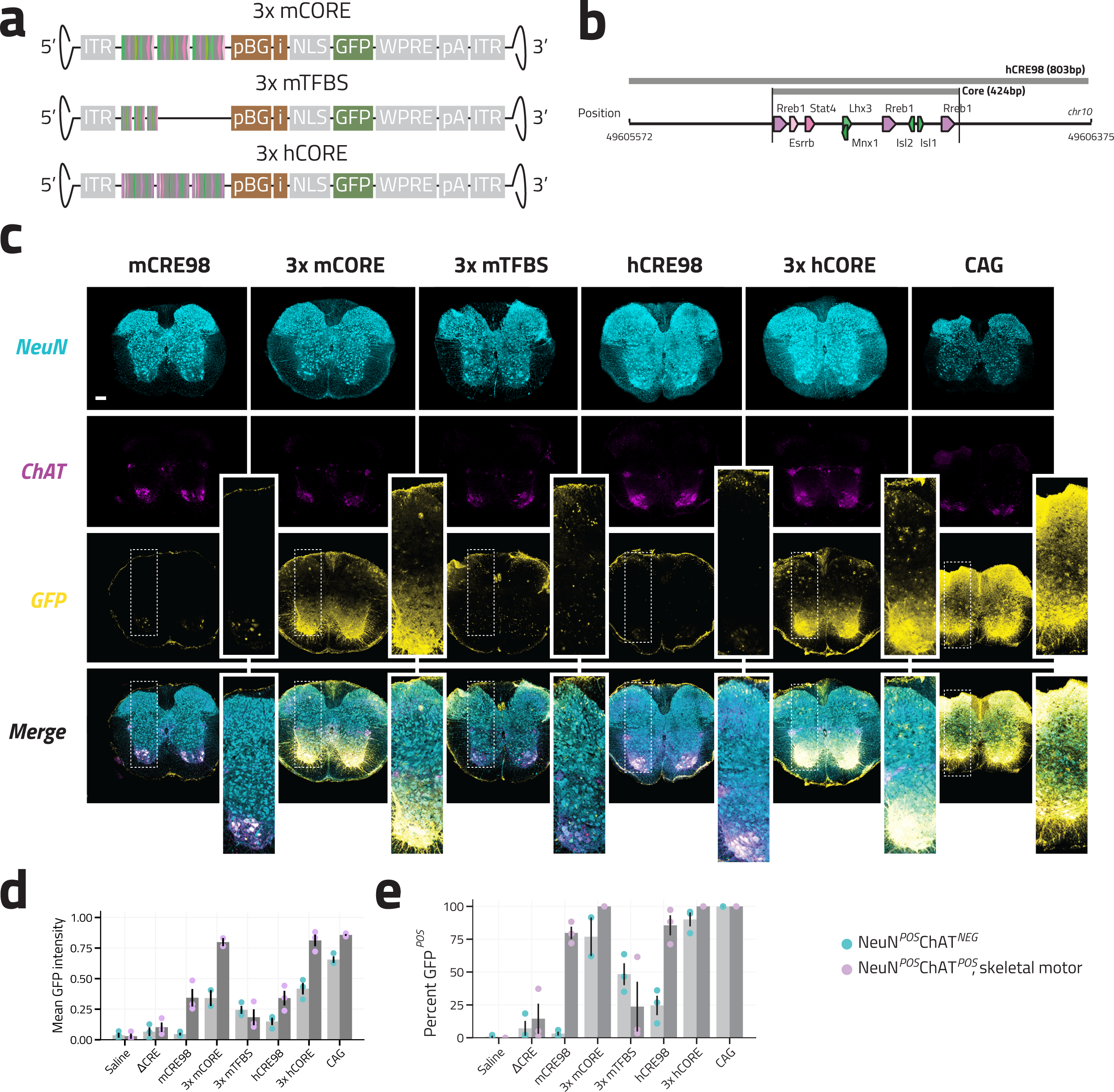
Evaluation of synthetic and human CRE98-derived constructs. **(a)** Synthetic AAV screening construct vector genome maps (not drawn to scale). ITR, inverted terminal repeats; pBG, minimal beta-globin promoter; i, intron; NLS, nuclear localization sequence; GFP, green fluorescent protein. WPRE, woodchuck hepatitis virus post-transcriptional response element; pA, poly-A tail. **(b)** Native genomic context of human CRE98 (hCRE98), showing TF-binding sites (TFBS) along genomic coordinates of chromosome 10. **(c)** Representative IHC images of T1-L4 spinal cord sections 14 days after P0 ICV injection of synthetic CRE reporter AAVs (NeuN, cyan; ChAT, magenta; GFP, yellow). mCRE98 and CAG images reproduced from Figure 3a. Scale bar 250 μm. **(d)** Quantification of mean GFP signal intensity and **(e)** percentage of GFP*^POS^*cells in off-target spinal neurons (NeuN*^POS^* ChAT*^NEG^*, blue) and on-target spinal motor neurons (NeuN*^POS^* ChAT*^NEG^*, purple) for saline-injected and ΔCRE controls (n = 3 animals each), CRE98 (n = 3 animals), synthetic constructs (n = 2-3 animals), and CAG (n=3 animals). Mean enrichment across all spinal cord sections per animal denoted by each point, mean across all animals per condition denoted by bar plot, standard deviation across all animals denoted by line. ΔCRE, mCRE98, and CAG data re-plotted from Figure 3.

Parallel testing of AAV reporter constructs incorporating either the full-length homologous human CRE98 sequence (hCRE98, 803 bp) or triplicate concatemer repeats of the homologous core TF-binding region (3hCORE) (**Figure 5b**) revealed similar trends to those observed with the corresponding murine sequences, with the 3hCORE construct driving increased expression compared to hCRE98, albeit with reduced selectivity (**Figures 5d, e**). Within the context of the mouse spinal cord, the human hCRE98 construct performed more poorly than the mouse CRE98 (mCRE98) element, as measured by on-target expression strength (0.34 ± 0.06), off-target expression strength (0.15 ± 0.03), and off-target hit rate (24.6 ± 7.3%). However, hCRE98 remained more MN-selective than any other CRE construct tested in the mouse spinal cord after mCRE98 and mouse CRE119, suggesting preservation of CRE98 function across mouse and human.

### CRE98 function is preserved in non-human primate

Cell-type-restricted CREs that preserve selective expression in viral contexts across diverse mammalian species have been previously reported (21, 23–25). Given the partial selectivity of hCRE98 in mouse, we sought to examine whether mCRE98 retains its spinal MN-selective expression in a non-human primate (NHP) context. To this end, wild-type adult, AAV9 neutralization antibody-naïve cynomolgus macaques (n = 3) were singly dosed with 1.0 x 10^13^ viral genome copies (1 mL) via intra cisterna magna (ICM) injection of a barcoded synthetic promoter AAV9 library that included CRE98-pBG- and mCORE-pBG-driven barcoded reporters (**Figures 6a, 6b)**. A CAG-promoter as well as the neuron-specific human *Synapsin 1* promoter (hSYN1), known to drive MN expression in the spinal cord in mice (50), were included in the screen as reference points for expression levels and specificity. Four weeks post-injection, tissues were dissected, and relative regulatory-element driven expression was quantified by bulk RNA-seq after normalization to dosing viral material (**Methods**).

**Figure 6.**
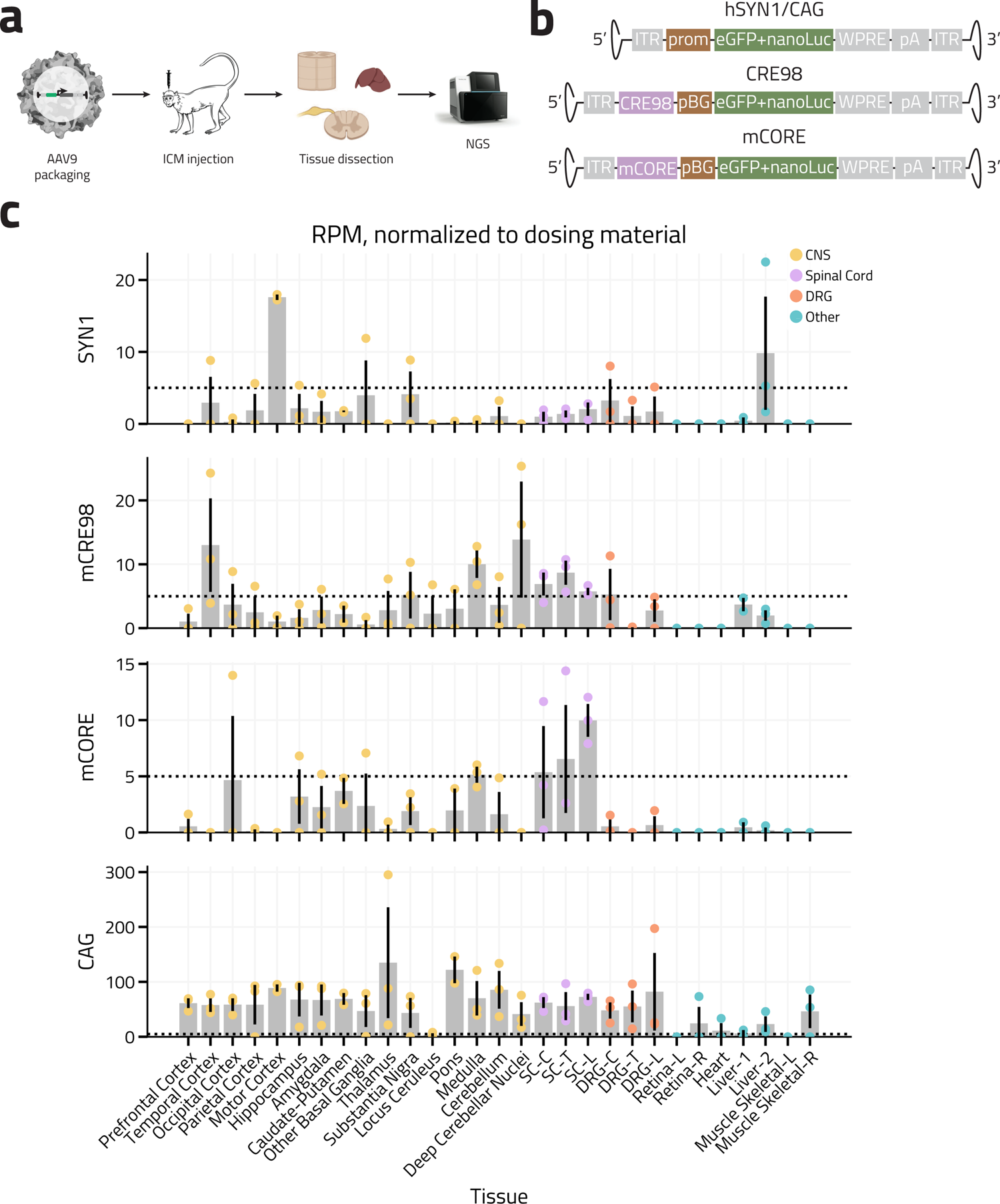
CRE98 drives spinal cord-selective expression in the macaque. **(a)** Experimental design for NHP study. NGS, next-generation sequencing. **(b)** AAV screening construct vector genome maps (not drawn to scale). ITR, inverted terminal repeats; prom, CAG or hSYN1 promoter; pBG, minimal beta-globin promoter; mCORE, mCRE98 core sequence; eGFP, enhanced green fluorescent protein; nanoLuc, luciferase reporter; WPRE, woodchuck hepatitis virus post-transcriptional response element; pA, poly-A tail. **(c)** Relative (normalized to dosing material) expression of hSYN1-, mouse CRE98 (mCRE98)-, mouse core CRE98 (mCORE)-, and CAG-driven AAV9 reporter constructs across tissues (on-target spinal cord in purple, off-target CNS-derived tissues in yellow, DRG in orange, and other in blue) harvested 28-days post-dosing from AAV9 neutralization-naïve cynomolgus monkeys (n = 3 animals). Normalized expression value of 5 denoted by dotted line across all constructs.

Relative to the hSYN1 promoter, mCRE98 and mCORE demonstrated significantly reduced off-target expression in motor cortex (17.5-fold and 4e3-fold decreased, respectively) and concomitantly increased expression in the spinal cord at all assessed levels (C4: 7.0-fold and 5.4-fold; T5: 6.3-fold and 4.8-fold; L1: 2.8-fold and 4.9-fold increased, respectively, **Figure 6c**). The occipital cortex and medulla exhibited the highest level of off-target CRE-driven expression for both constructs. All three constructs demonstrated low, though nonzero, levels of reporter expression in DRG, with the greatest relative specificity attained by the mCORE construct. Finally, the CAG construct drove approximately 10-fold greater reporter expression in the spinal cord than either CRE, with similarly high levels of off-target expression in DRG and across the CNS. Together, these data show that, within the context of AAV delivery of genetic material, the mouse CRE98 and its newly identified core functional region drive robust and selective payload expression in the macaque spinal cord relative to the canonical pan-neuronal hSYN1 promoter. Therefore CRE98, or the more compact CRE98 core sequence, represent potentially valuable tools capable of conferring MN-restricted transgene expression upon incorporation into recombinant AAV that merit further higher-order vertebrate testing for potential therapeutic purposes in humans.

## DISCUSSION

The identification and application of cis-regulatory elements to refine viral transgene expression profiles is a burgeoning field with substantial promise for both preclinical and clinical use. Within this domain, lower motor neurons of the spinal cord underlie several devastating degenerative disorders and represent an important investigational target for which no such CREs have been described to date, in part because of the difficulty of experimentally isolating this cell population. In this study, we transcriptionally and epigenetically profile skeletal motor neurons in mouse to identify two cis-regulatory elements, CRE98 and CRE119, that drive significant motor neuron-selective transgene expression in an AAV context. We further identify a 280 bp central core TF-binding region that underlies this MN-selective expression in mouse CRE98, demonstrate its conservation in humans, and find improved spinal cord expression compared to the pan-neuronal hSYN1 promoter in an NHP context.

CRE98 appears to be strongly conserved across multiple mammalian species, including mouse, rat, macaque, and human. This conservation highlights the functional importance of this element in regulating genes essential for cholinergic signal transduction and is further evidenced by preserved function of both the human CRE98 sequence in mouse and the mouse CRE98 in NHP. As many viral therapies fail in human testing despite promising preclinical animal studies, these findings provide an extra layer of reassurance arguing in favor of preserved CRE98 function for potential clinical applications.

Although our efforts were focused on CRE98 in light of its favorable expression profile, we also examined the sequence composition of CRE119. Only two motifs whose associated TFs demonstrated spinal MN-enriched expression in our RNA-seq dataset (albeit with low levels of expression) were identifiable within the CRE119 sequence: Tfcp2l1 (transcriptional regulator traditionally associated with embryonic stem cell pluripotency and known target of the MN-enriched Stat3 TF) and Nr2e1 (cell cycle regulator of neuronal stem cells important for CNS and retinal development (51, 52), **Supplementary Figures 4b, 4c**). Neither of these TFs is traditionally associated with MN development, and surprisingly, of the motifs identified in CRE98, only the Rreb1 motif was shared with CRE119. Furthermore, a developmental ATAC-seq dataset in mouse skeletal MNs demonstrated that the endogenous CRE98 sequence is chromatin accessible at all age ranges assessed (E10.5 to P2 years), whereas CRE119 exhibits delayed accessibility beginning at P13 (30). Together, these findings represent an exciting avenue of future exploration to understand MN development and maturation, as CRE98 and CRE119 appear to utilize distinct molecular mechanisms to achieve MN-selective control of gene expression across different developmental stages.

Despite its substantially improved specificity profile and increased expression in NHP spinal cord relative to CAG or hSYN1, CRE98-driven on-target expression remains substantially reduced relative to the ubiquitous CAG promoter (15.4- and 8.9-fold reduced in mouse and NHP, respectively, **Supplementary Figure 3f**, **Figure 6c**) currently employed in the majority of MN-focused gene therapies in human (3, 10–12, 53). These constructs may not achieve sufficient payload expression levels necessary for therapeutic effect in all protein replacement or rescue applications, however, they may represent promising avenues for approaches that favor selectivity over raw expression (including CRISPR/Cas9-based gene editing) or in conditions where supraphysiologic expression (i.e. CAG promoter-drive) engenders toxicity (14, 54–56). Additionally, our finding that CRE98 core element concatemers have higher on-target expression than the full-length sequence with partially preserved on-target enrichment relative to CAG suggests that additional synthetic sequence refinement may further optimize CRE98 for gene delivery applications, though this remains to be explored. Finally, the increased expression with concatemerization suggests that CRE98 inclusion does not deplete the pool of native TFs or alter baseline transcription to the detriment of normal cellular function (57), a frequent concern levied against the inclusion of highly-expressing viral gene therapies. Given widespread expression of ChAT in the CNS, it also remains to be explored to what degree CRE98 confers cholinergic selectivity in this important clinical context as well.

In summary, CRE98 and its compact core sequence represent valuable tools capable of conferring MN-selective transgene expression upon incorporation into recombinant AAV vectors. The identification of these elements and their preserved function in the NHP context reaffirm the strategy of epigenetic profiling for the identification of candidate elements, with substantial utility in developing scientific tools, unraveling endogenous mechanisms of restricted gene expression, and generating new therapies for human pathology.

## MATERIALS AND METHODS

### Mice

Animal experiments were approved by the National Institute Health and Harvard Medical School Institutional Animal Care and Use Committee, following ethical guidelines described in the US National Institutes of Health Guide for the Care and Use of Laboratory Animals. For INTACT, we crossed homozygous *Chat-IRES-Cre* (The Jackson Laboratory Stock # 006410) mice with homozygous *SUN1-2xsfGFP-6xMYC* (The Jackson Laboratory Stock # 021039) mice and used adult (6-12-week-old) heterozygous male and female F1 progeny. For CRE screening, we used adult (5-7-week-old) C57BL/6J (The Jackson Laboratory, Stock # 000664) mice. All mice were housed under a standard 12 hr light/dark cycle.

### INTACT nuclear isolation for sequencing

Nuclear isolation: Animals were deeply anesthetized with isoflurane (1-3% in air) and sacrificed by decapitation (surgical scissors) before dissection and processing of thoracic and lumbar spinal cords. Spinal cords from two mice were combined per experimental replicate. Spinal cords were dounced 10 times in 5 mL of HB buffer (0.25 M sucrose, 25 mM KCl, 5 mM MgCl_2_, 20 mM Tricine-KOH pH 7.8, 1 mM DTT, 0.15 mM spermine, 0.5 mM spermidine, DTT 1 mM [Sigma], half of a Roche proteinase inhibitor tablet) in a 7 mL dounce-homogenizer, supplemented with IGEPAL CA-630 to final concentration of 0.3%, and dounced an additional ten times. Nuclei were filtered through a 40 μm strainer and combined 1:1 with 50% iodixanol. Nuclei were then underlaid with a gradient of 1 mL 40% iodixanol and 1 mL 30% iodixanol and centrifuged at 10,000*g* for 19 minutes using the “no brake” setting. Purified nuclei (0.8 mL) were then collected from the 30/40% interface. A 20 μL aliquot was taken for visual inspection and cell counting, with typical yield of ∼1 million nuclei per two spinal cords. For RNA-seq experiments, all buffers were supplemented with 1.5 μL/mL RNAse inhibitor (Promega).

Affinity purification: A modified version of the published INTACT protocol (28) was applied, with the following changes. 1) Each experimental sample was split across two LoBind tubes to accommodate the larger volumes obtained. 2) Immunoprecipitation (IP) was completed over 30 minutes of end-to-end rotation at 4°C. 3) Flowthrough during the IP step was also collected, filtered through a 20 !m CellTRICs filter, and centrifuged at 800*g* for 10 minutes at 4°C before resuspension in 400 μL wash buffer (buffer HB supplemented with 0.4% IGEPAL CA-630) with careful pipetting. 4) The final immunoprecipitate was then resuspended 400 μL wash buffer. A 20 !L aliquot was taken from each sample for cell counting and verification of effective purification of GFP-positive (GFP*^POS^*) nuclei. Typical yield was approximately 25,000 GFP*^POS^* nuclei per spinal cord (>95% GFP*^POS^*) in the immunoprecipitate and approximately 150,000 nuclei per spinal cord from flowthrough (<1% GFP*^POS^*).

### ATAC-seq library preparation and sequencing

ATAC-seq was performed using the OMNI-ATAC protocol (58), starting with the steps that immediately follow cell lysis. Nuclei (50,000) were pelleted at 500 *g* for 10 minutes at 4°C in a fixed angle centrifuge and resuspended in 50 μL transposition buffer (prepared from Illumina kit components TD buffer and 100 nM transposase [Illumina], as well as 0.01% digitonin, 0.1% Tween-20) via careful pipetting. Transposition occurred at 37°C in a thermomixer at 1000 rpm for 30 minutes before resuspension in 250 μL of DNA binding buffer (Zymo DNA Clean and Concentrator Kit) and reaction cleanup per manufacture instructions, with final elution volume of 21 μL in elution buffer. Samples were amplified following NEBNext High-Fidelity 2X PCR Master Mix manufacturer protocols (NEB) in a total reaction volume of 50 μL (1.25 μM each of i5 and i7 sequencing primers) at 72°C for 5 minutes, [98°C for 10 seconds, 63°C for 30 seconds, and 72°C for 1 minute] x10 cycles. PCR samples were re-purified following the Zymo manufacturer protocol and eluted into 21 μL of elution buffer. Size selection was performed via agarose gel (150-1000 bp) followed by Qiagen MinElute gel extraction according to the manufacturer’s instructions, with two serial 10 μL elution steps. Seventy-five bp paired-end sequencing was performed on an Illumina NextSeq500.

### Sequencing, alignment, and genome browser track generation

All experiments were sequenced on the NextSeq 500 (Illumina). Seventy-five bp paired-end reads were obtained for all datasets. ATAC-seq experiments were sequenced to a minimum depth of 20 million reads. RNA-seq experiments were sequenced to a depth of at least 30 million reads. All samples were aligned to the mm10 genome using default parameters for the Subread alignment software (subread-1.4.6-p3) (59) after quality trimming with Trimmomatic v0.33 (60) with the following command: `java -jar trimmomatic-0.33.jar SE - threads 1 -phred33 [FASTQ_FILE] ILLUMINACLIP:[ADAPTER_FILE]:2:30:10 LEADING:5 TRAILING:5 SLIDINGWINDOW:4:20 MINLEN:45`. Truseq adapters were trimmed out in RNA-seq experiments; Nextera adapters were specified for ATAC-seq data. To generate UCSC genome browser tracks for ATAC-seq data, all aligned bam files for each replicate of a given experiment were pooled and converted to BED format with bedtools `bamtobed`. The 75 bp base reads were extended in the 3’ direction to 200 bp (average fragment length for ATAC-seq experiments as measured by bioAnalyzer) with the bedtools `slop` command using the following parameters: `-l 0 -r 125 -s`. Published mm10 ChIP-seq blacklisted regions (38) were filtered out using the following command: `bedops –not-element-of 1 [BLACKLIST_BED_FILEPATH]`.

The filtered BED files were converted to coverageBED format using the bedtools `genomecov` command with the following options: `-scale [NORM_FACTOR to scale each library to 20M reads] –bg`. Finally, `bedGraphToBigWig` (UCSC-tools) was used to generate the bigWIG files displayed on browser tracks throughout the manuscript.

### ATAC-seq peak calling and quantification

Regions of ATAC-seq enrichment were determined using MACS2 (v 2.1.0) parameters `-p 1e-5–nolambda –keep-dup all –slocal 10000`, as previously described (37). Peaks from individual sample replicates were intersected to find only reproducible regions of enrichment. Blacklist regions were removed as previously described (38). To identify sites with enriched ATAC-seq signal (peaks), we applied the IDR pipeline (61) using the MACS2 peak calling algorithm (62) with the following parameters: `–nomodel –extsize 200 –keep-dup all`. An IDR threshold of 0.01 was used for self-consistency, true replicate, and pooled-consistency analyses. The ‘`optThresh`’ cutoff was then used to obtain a final set of high-confidence, reproducible ATAC-seq peaks for each sample.

To produce a final list of reference coordinates containing 42,680 genomic regions that were accessible in at least one spinal ATAC-seq sample, the MACS2 peaks for each experimental replicate were unioned using the `bedops --everything` command. Bedtools `merge` was then used to combine any peaks that overlapped in this unioned bed file; in this way, any region that was significantly called a peak in at least one ATAC-seq dataset was incorporated in the final aggregated peak list. The `featureCounts` package was then used to obtain ATAC-seq read counts for each of these accessible putative CREs for downstream enrichment analyses (63).

### CRE conservation analysis

Average conservation scores for all putative spinal cord CRE sequences across the placental clade were calculated using the `bigWigAverageOverBed` tool, inputting PhyloP scores of each sequence derived from the publicly available `mm10.60way.phyloP60wayPlacental.bw` dataset (http://hgdownload.cse.ucsc.edu/goldenpath/mm10/phyloP60way/) (64, 65). After plotting the conservation score, we determined the conservation score of the 95^th^ percentile of this distribution (PhyloP score = 0.5) and chose it as a minimal conservation score needed to classify any CRE as conserved. Using this cutoff, 20 CREs were classified as conserved and included for subsequent selection of CREs for testing.

### Motif identification

To identify putative transcription factor-binding sites, the FIMO MEME analysis tool (Find Individual Motif Occurrences, v5.3.3 (49)) was applied, scanning for motifs contained in the JASPAR 2020 transcription factor motif database (66) using the following command: `fimo –oc . –verbosity 1 –thresh 1.0E-4 JASPAR2020_CORE_vertebrates_redundant_pfms_meme.txt [INPUT_FILE]`, where INPUT_FILE denotes the full mouse CRE98 and CRE119 sequences. To identify functional sites, the output motif list was filtered for TFs that were selectively expressed in INTACT-purified ChAT*^POS^* cells of the spinal cord relative to ChAT*^NEG^*depleted flow-through (defined as FC ≥ 2.0, q < 0.05) by RNA-seq.

### RNA purification, library preparation, and sequencing

Nuclei from INTACT immunopurification were immediately pelleted at 500 *g* for 10 minutes at 4 °C in a fixed angle centrifuge and resuspended in 350 μL RLT buffer [Qiagen]. RNA purification was performed following the Qiagen RNEasy MinElute manufacturer’s protocol, with a final elution volume of 20 μL. Libraries were prepared following the SMART-seq manufacturer’s protocol [Takara Bio] with 15 amplification cycles and purified following the AMPure XP [Beckmann Coulter] manufacturer’s instructions.

### RNA-seq alignment and quantification

To quantify gene expression, we applied the `featureCounts` package (63) after aligning RNA-seq libraries to the genome as described above, using a custom filtered annotation file: `gencode.v17.annotation.gtf filtered for feature_type=“gene”, gene_type=“protein_coding” and gene_status=“KNOWN”` to obtain gene counts for each sample. These counts tables were TMM-normalized using the `EdgeR` software analysis package (67). Any genes that were not expressed in at least 3 samples with TMM-normalized counts per million (CPM) >1 were dropped from further analysis. Differential expression (DE) analyses were conducted using the `voom/limma` analysis software packages (requiring FDR-corrected q<0.05) (68, 69).

### Amplification of CREs

The same constructs and cloning protocol applied in Hrvatin et al. 2019 (22) were used to generate viral constructs in this manuscript., albeit with an NLS-eGFP transgene instead of GFP. PCR primers were designed using primer3 2.3.7 (69) such that a 150–400 bp flanking sequence was added to each side of the CRE. The forward primers contained a 5’ overhang sequence for downstream in-Fusion [Clonetech] cloning into the AAV vector (5’-GCCGCACGCGTTTAAT). The reverse primers contained a 5’ overhang sequence containing the recognition sites for AsiSI and SalI restriction enzymes (5’-GCGATCGCTTGTCGAC). Hot Start High-Fidelity Q5 polymerase [NEB] was used according to manufacturer’s protocol with mouse genomic DNA as template. The following primer sequences were used for the CREs evaluated in this study:

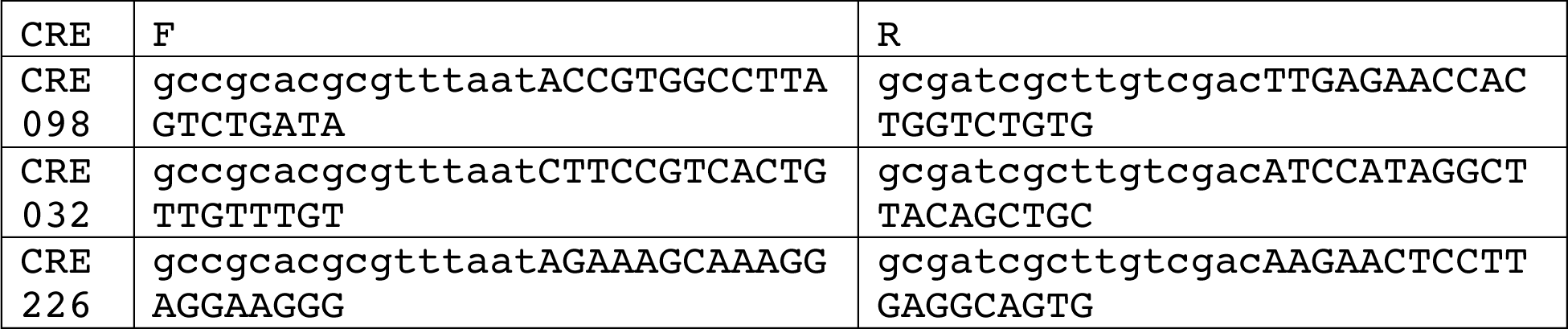
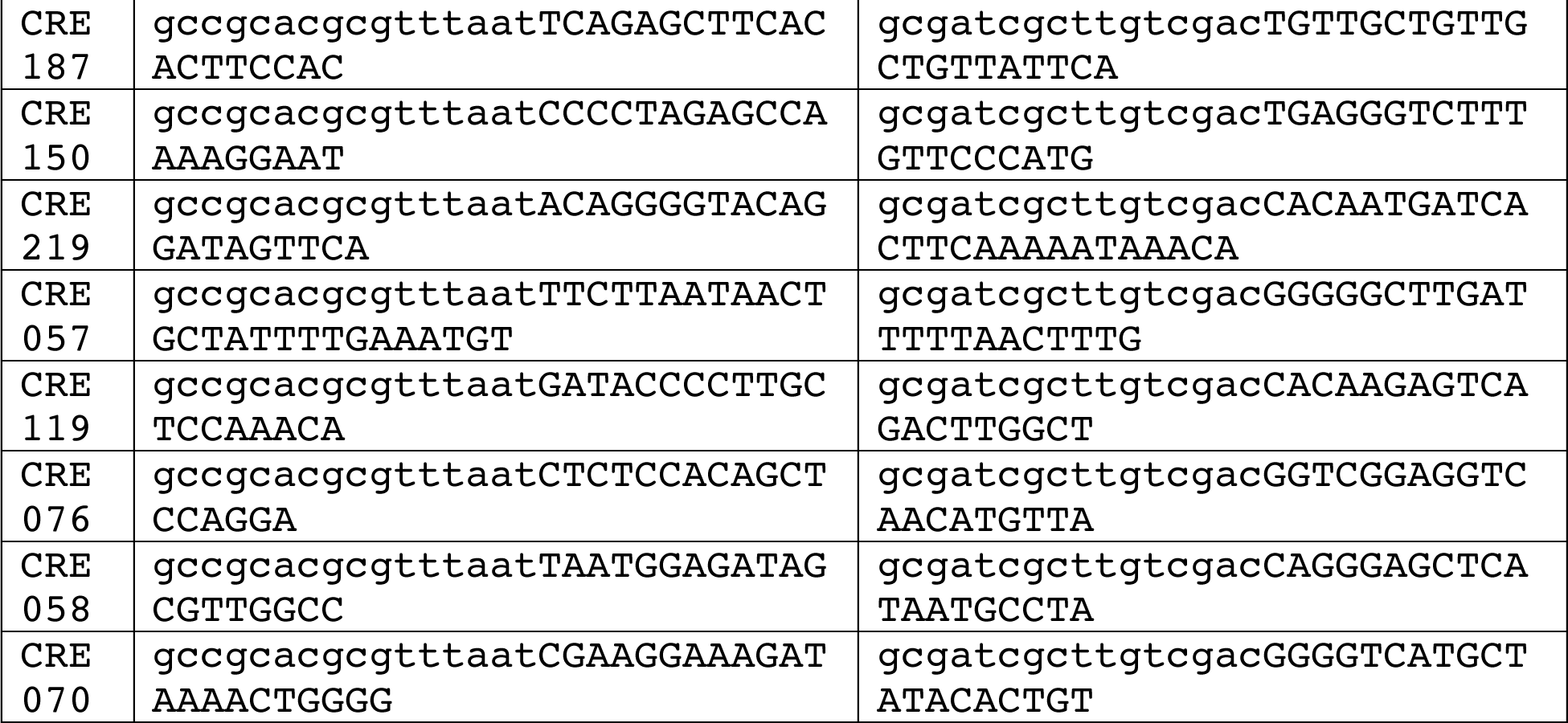

The unpurified PCR products from the genomic PCR were used as templates for the barcoding PCR. A forward primer containing the sequence for downstream in-Fusion [Clonetech] cloning into the AAV vector (5’-CTGCGGCCGCACGCGTTTA) was used in all reactions. Reverse primers were constructed featuring (in the 5’ → 3’direction): 1) a sequence for downstream in-Fusion [Clonetech] cloning into the AAV vector (5’-GCCGCTATCACAGATCTCTCGA), 2) a unique 10-base barcode sequence, and 3) sequence complementary with the AsiSI and SalI restriction enzyme recognition sites that were introduced during the first PCR (5’-GCGATCGCTTGTCGAC). Three different reverse primers were used for each of the GREs amplified during the genomic PCR. Hot Start High-Fidelity Q5 polymerase [NEB] was used according to the manufacturer’s protocol.

The CAG promoter was digested from Vector Biolabs plasmid 7075 using MluI/AsiSI restriction and cloned into the viral construct by ligation. The ΔCRE construct was similarly constructed by ligation after MluI/AsiSI restriction to remove the CRE from the viral vector construct.

For enhancer truncation experiments, the following primers were used to clone portions of the full-length CRE98 construct from mouse genomic DNA, which were then inserted into the final vector construct as described previously:

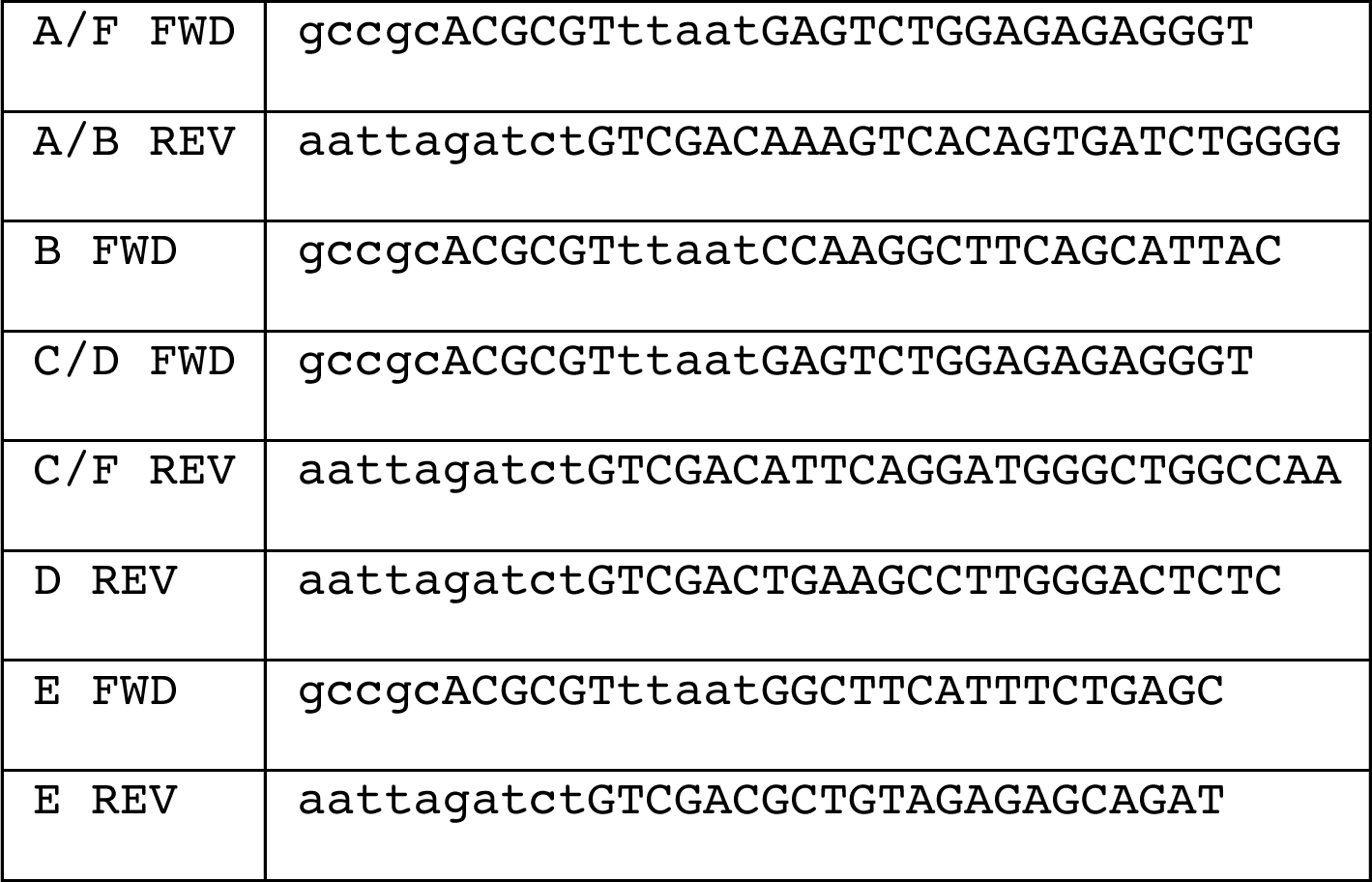

The 5KO cloning template was ordered as a gBlock [IDT] with the following sequence and cloned into the viral construct as described above with primers A/F FWD and C/F REV: 5’ – GAGTCTGGAGAGAGGGTGGGAGCAGCCATTCTGCAGCAGTGCCTTCTTGGGGTCATGGGTCTGT AGGTGCTGCTGTGGAGGGAGAGATCAGCCTATTCTGGCTTCATTTCTGAGCTGCAAACTGCCTG GGTGTCTGGAGAAGCAGGTTGGCGTGGTGGTTAGCAGTGCGTGGGCGGGGTTGCCCGCTCTTGA TTTATGATTTCTTTGTCTCTGTGGAAAGGCTTTAGTTCCAATGACACTCAGGAGCCTCTGGATT CCAGTAGAACGTTCTCAGGCCTCACCAACCCCTCCCCTGTGTGCTGCCTTTGGGAGAGTCCCAA GGCTTCAGCATTGCCTCTACTGCTACATAGGCTCAGATTCAAAAGAACAGAGTGGCCCACGTCA AGTCTGATGGCTGGAAGCCAGAGGACTATGTGTCTGCCTTGCGCCCATCCTGAATGCCCAGACT CGGACAATGGAGTAGGTACAGAAGGGTAAAGACAGTGTCTTCTGTACCAGTAAGTGGGCCCTGA TCTGCTCTCTACAGCTTCCAGAGAAAGGGCCTGGCCAATGAGCGGCCTTTTGAGTAGCAGATAC CTCACATGCATTCTGATAGAAAGCCTGGCCCCAGATCACTGTGACTTT – 3’

### AAV preparation for murine injection

Individual AAV constructs (100 μg) were packaged into AAV.PHP.eB (42) at the Boston Children’s Hospital Viral Core. The titers (2–20 × 10^13^ genome copies/mL) were determined by qPCR and normalized to 4 x 10^13^ genome copies/mL.

### Intracerebroventricular injection in mice

P0 mice were anesthetized with isoflurane (1-3% in air) and chilled on anodized aluminum blocks resting in ice prior to injection. Virus (4 μL) with 0.04% FastGreen dye [EMS, cat. 26364-05] was injected into the right lateral ventricle 0.8-1 mm laterally from the sagittal suture halfway between lambda and bregma (visible through the skin) via a 33-gauge 1701RN syringe [Hamilton, small hub RN NDL, custom length 0.5 inches, point 4 (45°), Cat. 7803-05]. Successful injection was confirmed by spread of dye into the third and contralateral lateral ventricles. Pups were then rewarmed in a 98°C incubator before they were returned to their home cages.

### Spinal cord, DRG, brain, and liver dissections for imaging

Anesthetized mice were transcardially perfused with 4% PFA followed by PBS. The relevant organs were dissected out and post-fixed with 4% PFA for 1-3 days at 4°C. Tissues were rinsed in 0.1 M phosphate buffer three times and set in agarose gel for mounting on the vibratome [Leica VT1000S]. Tissues were axially sectioned into ∼50 μm slices and transferred to 24-well plates for immunostaining.

### Immunostaining

Samples were washed in 50% ethanol solution to increase antibody permeability, then washed three times with PBS containing 0.3% TritonX-100 (PBST) and blocked for 1 hour at room temperature with PBST containing 10% donkey serum. Sections were incubated 24-48 hours at 4°C with primary antibodies, washed again four times in PBST, and incubated for 1.5 hours at room temperature with the appropriate secondary antibody (see below). After washing in PBST and PBS, samples were stained in PBS containing DAPI and mounted onto glass slides using anti-fade medium (Glycerol:PBS, 3:7).

The following primary antibodies were used: GFP (Chicken, Aves Labs Inc. GFP1020, 1:500), ChAT (Rabbit, Abcam ab181023, 1:100), NeuN (Mouse, Thermo Fisher Scientific MAB377MI, 1:500). The following secondary antibodies were used: Jackson ImmunoResearch Laboratories 703-545-155, Abcam ab150110, Life Technologies A32860.

### Imaging

Sections were imaged on a Leica TCS SPE confocal microscope using an ACS APO 20x/0.30 CS objective (Harvard NeuroDiscovery Center). Tiled areas of ∼1.2 mm by 0.5 mm were imaged at a single optical section to avoid counting the same cell across multiple optical sections. Channels were imaged sequentially to avoid optical crosstalk.

### Image quantification

For Figure 2, images were manually split into ventral and dorsal sections via freehand mask selection, and GFP intensity was measured across anatomical regions using the MeasureObjectIntensity module in CellProfiler [v4.2.4].

For all other spinal cord images, to identify the threshold intensity used to classify DAPI*^POS^* nuclei, we used three class thresholding where pixels in the middle intensity are assigned to background using the `IdentifyPrimaryObjects` module (Otsu thresholding method, adaptive strategy, typical diameter 10-100 pixels, threshold smoothing scale 1.3488, threshold correction factor 2, size of adaptive window 50 pixels) in `Cellprofiler` [v4.2.4]. NeuN*^POS^* objects were identified with the same method. ChAT*^POS^* objects were identified manually and by anatomical location (spinal MN in ventral horn, visceral MNs in lateral horn, and ChAT*^POS^* cells around central canal) using freehand mask selection. DAPI*^POS^* nuclei were classified as NeuN*^POS^* or ChAT*^POS^* by masking overlap with DAPI, NeuN, and ChAT objects. NeuN, ChAT, and GFP channel intensity was measured over the size of the DAPI mask.

To identify the threshold intensity used to classify each DAPI*^POS^* nucleus as either NeuN*^POS^* or ChAT*^POS^*, we first determined the background signal in the channel representing NeuN or ChAT by selecting multiple points throughout the area visually identified as background. These background points were masked as small circular areas (radius = 5.7 μm), over which the mean background signal was quantified. The highest mean background signal for NeuN or ChAT was conservatively chosen as the threshold for classifying DAPI*^POS^* cells as NeuN*^POS^* or ChAT*^POS^*, respectively, using the mean signal quantification across the DAPI*^POS^* mask obtained as described in the preceding paragraph.

To identify the threshold intensity used to classify each cell as GFP*^POS^*, we determined the GFP*^POS^* signal overlying all DAPI*^POS^* nuclei in the control saline condition as described above, and identified the 99th percentile of GFP expression as the threshold (**Supplemental Figure 3c**) for positivity. As CAG-driven GFP signal intensity was an order of magnitude greater than the CRE-driven constructs, two separate image acquisition settings were used to capture the full dynamic range of CRE- and CAG-driven expression separately (**Supplemental Figures 3c, 3h**).

For DRG, images were batch processed in CellProfiler [v4.2.54] to segment DAPI*^POS^* nuclei and quantify GFP expression using these respective channels. We identified nuclei based on shape and signal intensity using the `IdentifyPrimaryObjects` module. We set an adaptive threshold strategy using the Otsu method, adaptive window size 50 pixels, creating a mask over objects between 10-70 μm in diameter and filtered nuclei by mean intensity (<0.5), area (between 100-600 pixels), and eccentricity (maximum 0.8). We quantified GFP expression within the area of DAPI*^POS^* masks using the `MeasureObjectIntensity` module as described above.

## Nonhuman primate experiments

### AAV construction and production

The mCRE98 and mCORE plasmids were cloned at Biogen using a proprietary proviral vector backbone with unique barcodes. AAV9-mCRE98-pBG-eGFP-WPRE-polyA (P105) and AAV9-mCORE-pBG-eGFP-WPRE-polyA (P106) were individually produced at UMASS Medical School Viral Vector Core (Worcester, MA, USA), together with 5 control vectors and AAV9-hSYN11-eGFP-WPRE-polyA. Viral titers were quantified by ddPCR using a WPRE-specific primer/probe set. Equal copies of titrated vectors were spiked into a pooled barcoded library, which was individually transfected but purified in minipool fashion. The combined library, with total 220 unique vectors, was AKTA-purified [Cytiva] and resuspended in artificial cerebrospinal fluid buffer. The absence of promoter or reporter gene mutations and inclusion of all candidates with balanced pooling were confirmed by NGS.

### Non-human primate (NHP) dosing and tissue collection

Three AAV9 neutralization antibody-naïve cynomolgus monkeys were dosed with 1 mL of pooled library at the dose of 1.0 x 10^13^ vg/animal via a bolus ICM injection by Charles River Laboratory by following established procedures in compliance with IACUC protocols. Monkeys were euthanized 28-days post-dosing, and tissue samples were collected from 16 different brain regions, multiple sections of spinal cord, dorsal root ganglion, and other peripheral organs for biodistribution and vector expression analysis.

### Gene expression in NHP samples by amplicon-seq

Collected NHP tissues were weighted, homogenized in individual tubes in RLT lysis buffer [Qiagen, 79216] with beta-mercaptoethanol using Geno grinder. Homogenates were arrayed into a deep 96-well plate and subjected to total RNA extraction by mixing 180 μL of homogenates with 1 mL of Qiazol [Qiagen, 79306] and 200 μL of chloroform. The collected aqueous phase (250 μL) was subjected to RNA extraction by following a modified MagMAX mirVana Total RNA Isolation [Thermo Scientific, A27828] protocol by KingFisher Flex, with DNase I treatment included. Extracted RNA was quantified by Nanodrop, treated with eZDnase [Invitrogen 11766051] to remove trace amounts of genomic DNA contamination, followed by reverse transcription into cDNA in the presence or absence (negative control) of SuperScript IV [Invitrogen 18091050]. To estimate amplification cycles for different tissues, SYBR Green qPCR amplification was performed with Kapa HIFI HotStart [Roche, KK2602] using adapted primers across the barcode region. Amplicons for motor cortex, spinal cord, and DRG were generated under the same conditions, except the use of different amplification cycles. Dosing material was also amplified for normalization purposes. Amplicons were purified and confirmed using Agilent TapeStation. Samples were then indexed and sequenced 101bp paired-end on a NovaSeq S1 flowcell by Biogen’s NGS lab following established sequencing protocols.

## Acknowledgments

This work was supported by a sponsored research award from Biogen (M.E.G.), and NIH T32GM007753 (M.A.N.). The authors would like to acknowledge Jiaochen Shen for her support on cloning the CRM vectors used in the NHP study, and Thomas Carlile, Soumya Negi and Jeron Chen for their assistance with the NGS and analysis of the NHP specimens.

## Competing interests

M.A.N., E.C.G., M.E.G. and S.H. are inventors on a patent related to this work. M.A.N., E.C.G. and S.H. serve or previously served as consultants to Apertura Gene Therapy. M.E.G. received sponsored research support from Biogen and Apertura Gene Therapy. S.H. holds equity in Apertura Gene Therapy and was a Visiting Scientist at Biogen at the time of this work. KW was an employee of and holds equity in Apertura Gene Therapy. S.P.G., E.R., L.M., V.P. and S.D. declare no competing interests. B.L., X.L., A.D., S.C.L., R.K., C.H., and J.S. were employed by Biogen and hold shares.

---

**Supplementary Figure 1.**
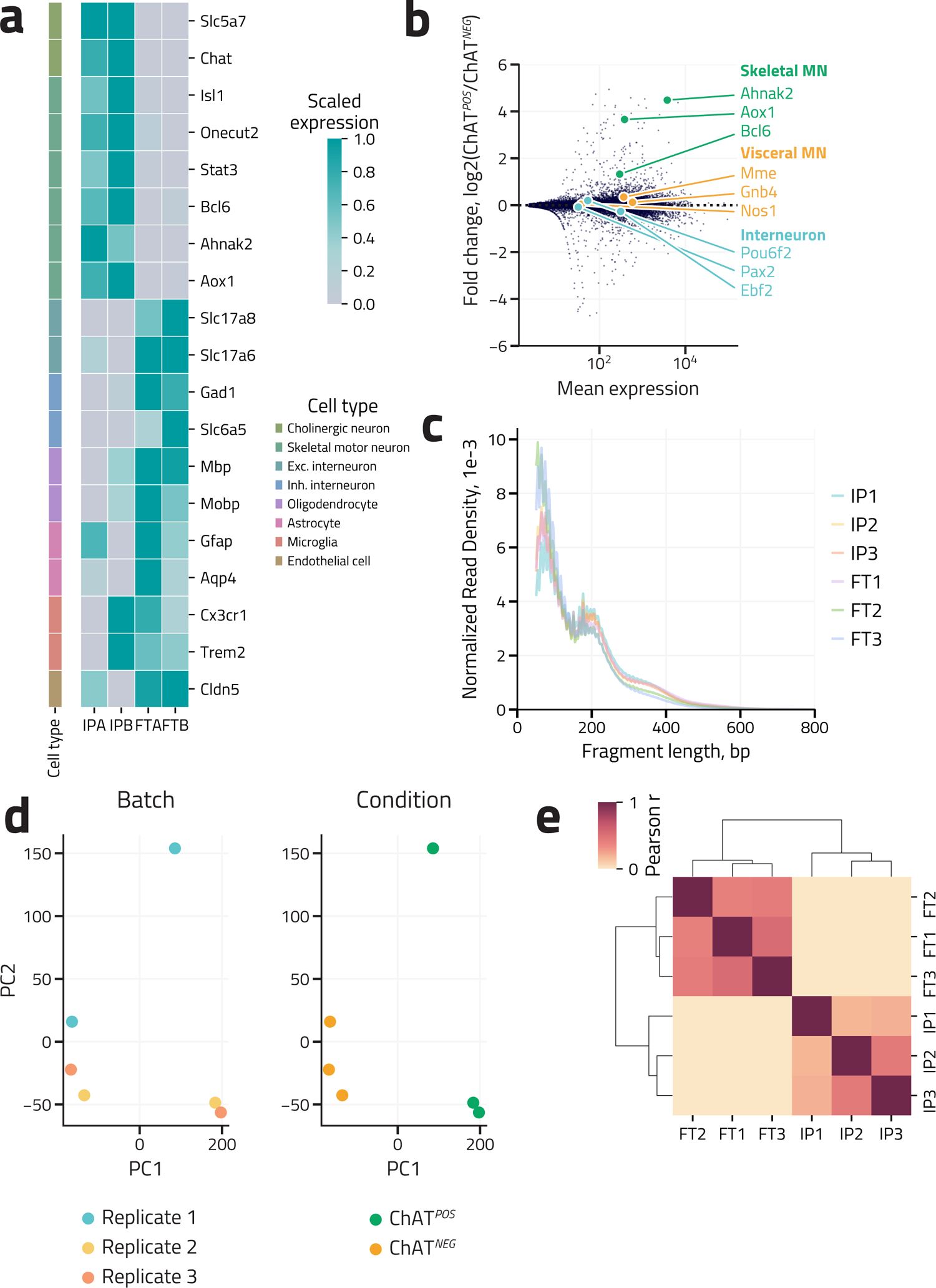
RNA- and ATAC-seq of immunopurified spinal motor neurons. **(a)** Heatmap of relative expression (scaled independently by maximum expression across four samples for each gene) of cell-type-specific markers across two bioreplicates (each bioreplicate comprises combined spinal cord tissue from two animals) of immunopurified spinal MN ChAT*^POS^* (IPA and IPB) and flowthrough ChAT*^NEG^* (FTA and FTB) nuclei. Cell types identified by literature review of marker genes (left, legend on right). **(b)** MA plot of MN-enrichment (log2[ChAT*^POS^*/ChAT*^NEG^*RNA-seq signal]) as a function of mean expression across all bioreplicates (each bioreplicate comprises combined spinal cord tissue from two animals) for all genes. Cholinergic neuron subtype-specific marker genes highlighted and labeled by class (skeletal MN in green, visceral MN in yellow, cholinergic interneuron in blue). **(c)** Fragment length distribution for all ATAC-seq samples from on-target ChAT*^POS^* spinal MN immunoprecipitate (IP) and off-target flowthrough (FT) bioreplicates (each bioreplicate comprises combined spinal cord tissue from two mice, n = 3 bioreplicates for each condition). **(d)** Principal component scatter plots of PC1 and PC2 for all ATAC-seq samples from on-target spinal MN ChAT*^POS^* immunoprecipitate (IP) and off-target ChAT*^NEG^* flowthrough (FT) bioreplicates (each bioreplicate comprises combined spinal cord tissue from two animals, n = 3 bioreplicates for each condition). Data are plotted twice to demonstrate PC1 driven by condition (right) rather than batch (left). **(e)** Pearson correlation heatmap of ATAC-seq bioreplicates for all ATAC-seq samples from on-target spinal MN ChAT*^POS^* immunoprecipitate (IP) and off-target flowthrough (FT) replicates. Bioreplicates are hierarchically clustered by the Pearson *R^2^*distance metric.

**Supplementary Figure 2.**
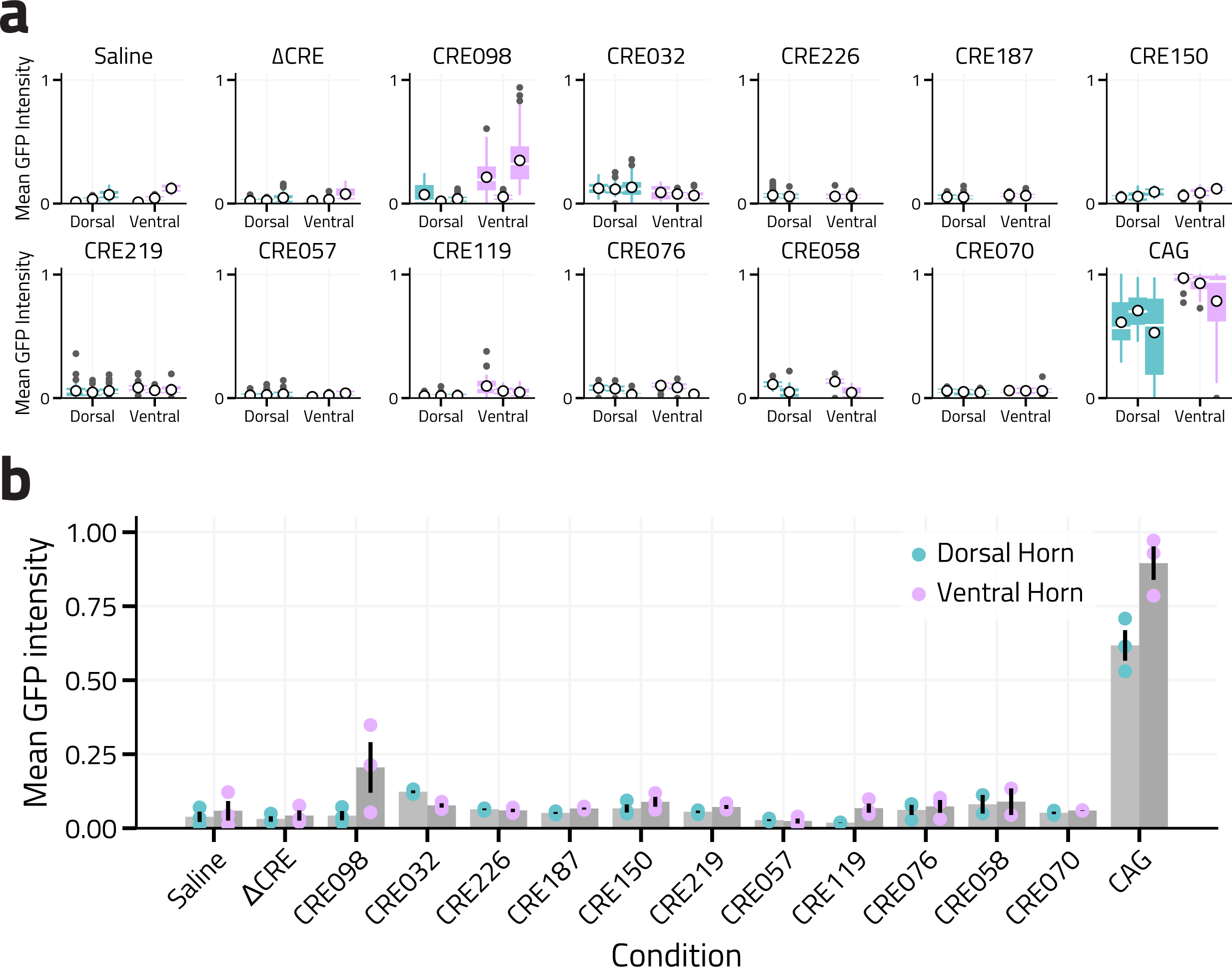
Quantification of GFP signal intensity in initial candidate CRE screening by confocal microscopy. **(a)** Quantification of GFP signal intensity in all off-target dorsal (blue) and on-target ventral (purple) spinal neurons as described in Figure 2. Individual cells are plotted separated by animal (x-axis labels, n = 2 for CRE226, CRE187, and CRE058; otherwise n = 3). Mean intensity across all cells per spinal cord section denoted by white circle, distribution of all cells by animal denoted by box-and-whisker plot (median denoted by white line, first and fourth quartiles by vertical line, outliers by gray points). (**b)** Raw quantification of mean GFP signal intensity averaged across all sections per animal for off-target dorsal horn (blue) and on-target ventral (purple) horn neurons used to calculate ventral enrichment in Figure 2e. Mean intensity across all sections per animal denoted by bar plot, standard deviation across animals denoted by vertical line. All images acquired and raw fluorescence intensity quantified at pBG promoter-optimized acquisition parameters.

**Supplementary Figure 3.**
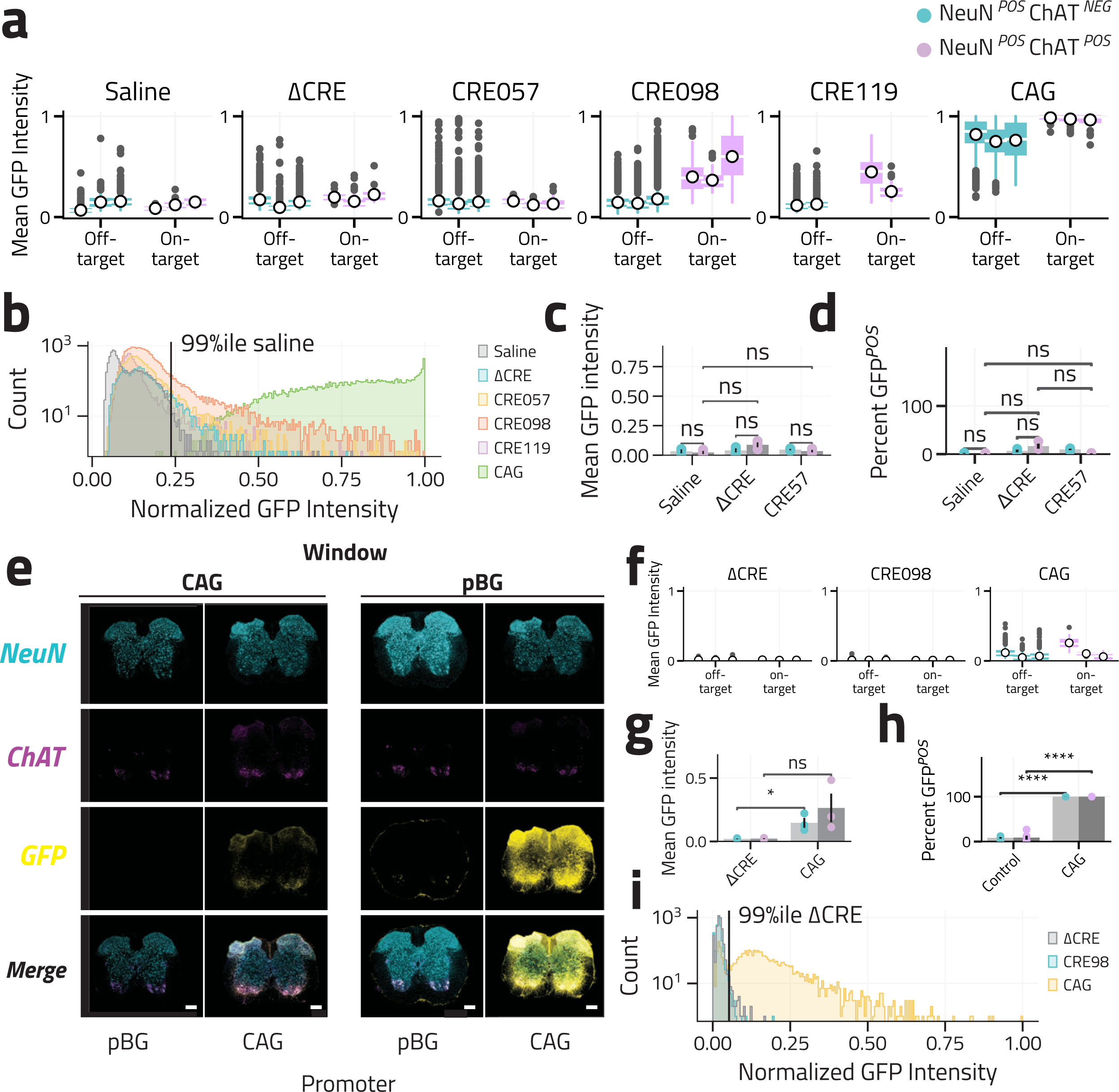
Quantification of CRE specificity by immunohistochemistry in spinal cord and DRG. **(a)** Quantification of GFP signal intensity in all off-target NeuN*^POS^*ChAT*^NEG^* (blue) and on-target NeuN*^POS^* ChAT*^NEG^* (purple) spinal neurons as described in Figure 3. Individual cells are plotted separated by animal (x-axis labels, CRE119 n = 2, all others n = 3). Mean intensity across all cells per spinal cord section denoted by white circle, distribution of all cells per animal denoted by box-and-whisker plot (median denoted by white line, first and fourth quartiles by vertical line, outliers by black points). **(b)** Histogram of GFP signal intensity across all nuclei at pBG promoter-optimized acquisition parameters settings used to acquire and quantify data shown in Figure 3 and Supplementary Figure 3a. Vertical black line indicates the 99^th^ percentile saline signal threshold used to categorize cells as GFP*^POS^*in Figure 3. **(c)** Quantification of mean GFP signal intensity and **(d)** percentage GFP*^POS^* in off-target (NeuN*^POS^* ChAT*^NEG^*, blue) and on-target spinal motor neurons (NeuN*^POS^* ChAT*^NEG^*, purple) for saline (n = 2), ΔCRE (n = 3), and CRE57 controls (n = 3) as described in Figure 3. Mean enrichment across all spinal cord sections per animal denoted by each point, mean across all animals per condition denoted by bar plot, standard deviation across all animals denoted by line. Images acquired and quantified at pBG promoter-optimized acquisition parameters for IHC signal as described in Supplemental Figure 3c. **(e)** Representative IHC images (NeuN, cyan; ChAT, magenta; GFP, yellow) of T1-L4 spinal cord sections 14 days after P0 ICV injection of CRE98 and CAG AAV constructs (construct denoted by top label) from animals as described in Figure 3 but with acquisition parameters optimized for CAG promoter-driven constructs. Images obtained at pBG promoter-optimized acquisition parameters reproduced directly from Figure 3A. Scale bar 250 μm. **(f)** Quantification of GFP signal intensity in all off-target NeuN*^POS^* ChAT*^POS^*and on-target spinal motor neurons as described in Figure 3 but plotted with CAG promoter-optimized acquisition parameters. Individual cells are plotted separated by animal for ΔCRE, CRE98, and CAG (n = 3 for all conditions). Mean intensity across all cells per spinal cord section denoted by white circle, distribution of all cells by section denoted by box-and-whisker plot (median denoted by white line, first and fourth quartiles by vertical line, outliers by black points). **(g)** Quantification of mean GFP signal intensity and **(h)** percentage of GFP*^POS^* cells in off-target spinal neurons (NeuN*^POS^* ChAT*^NEG^*, blue) and on-target spinal motor neurons (NeuN*^POS^* ChAT*^NEG^*, purple) for ΔCRE (n=3) and CAG constructs (n=3) as described in Figure 3, but with CAG promoter-optimized acquisition parameters. Mean enrichment across all spinal cord sections per animal denoted by each point, mean across all animals per condition denoted by bar plot, standard deviation across all animals denoted by line. **(i)** Histogram of GFP signal intensity across all nuclei at CAG-optimized acquisition parameters. Vertical black line indicates the 99^th^ percentile ΔCRE control signal threshold used to categorize cells as GFP*^POS^* in the CAG promoter-optimized acquisition parameters. Significance measured by Bonferroni-corrected two-tailed t test. *** q < 0.001, ** q < 0.01, * q < 0.05, *ns* q > 0.05.

**Supplementary Figure 4.**
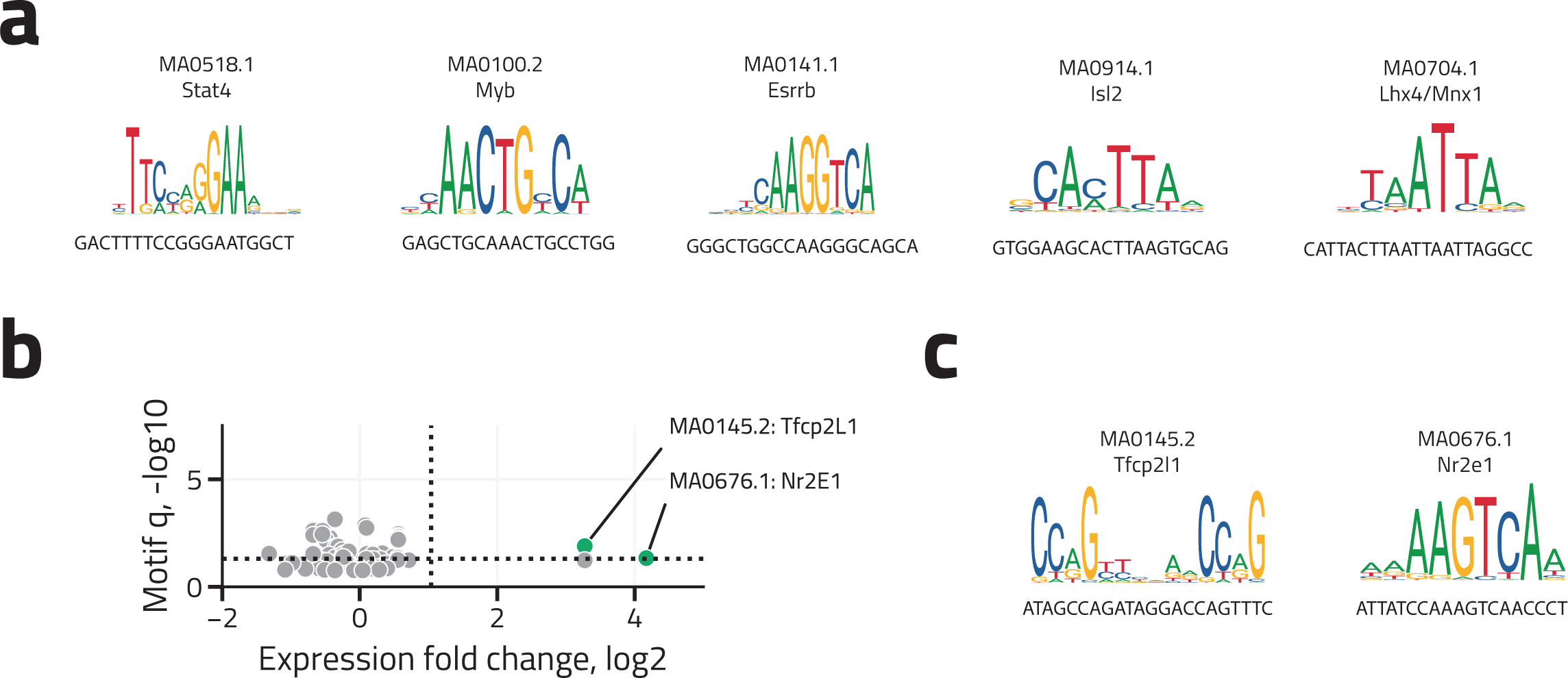
Truncation and motif analyses of CRE98 and CRE119. **(a)** Position weight matrix representation for significantly enriched CRE98 TFBS motifs (defined as > 2-fold expression enrichment in MNs by RNA-seq, FDR-corrected q < 0.01, and motif-enrichment q < 0.01) are shown with associated native mouse genomic sequence. **(b)** Scatter plot of significance (-log10 [Benjamini-Hochberg corrected TF motif enrichment q-value in CRE119]) as a function of MN-enrichment (log2[ChAT*^POS^*/ChAT*^NEG^*RNA-seq signal]). Motif enrichment q-values derived from FIMO. Motifs that exhibit > 2-fold expression enrichment in MNs (vertical dotted line), FDR-corrected q < 0.01 (horizontal dotted line), and motif-enrichment q < 0.01 are denoted in green and labeled by JASPAR motif ID and associated transcription factor(s). **(c)** Position weight matrix representation for significantly enriched CRE119 TFBS motifs (defined as > 2-fold expression enrichment in MNs by RNA-seq, FDR-corrected q < 0.01, and motif-enrichment q < 0.01) are shown with associated native mouse genomic sequence.

**Supplementary Figure 5.**
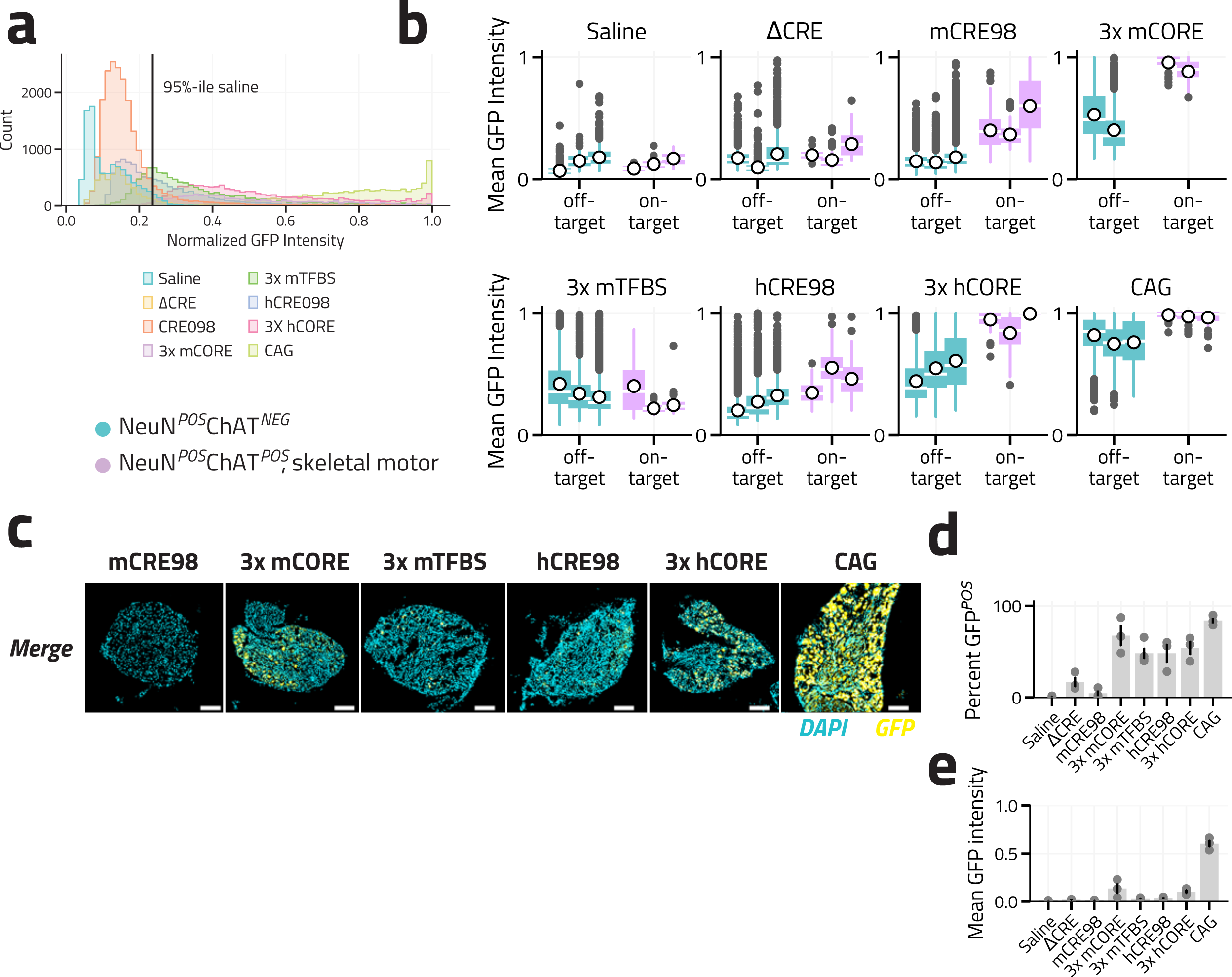
Quantification of synthetic and human CRE98-derived constructs in spinal cord and DRG. **(a)** Histogram of GFP signal intensity across all nuclei for synthetic and human constructs at pBG promoter-optimized acquisition parameters as described in Figure 5. Vertical black line indicates the 99^th^ percentile saline control signal threshold used to categorize cells as GFP*^POS^*. **(b)** Quantification of GFP signal intensity in all off-target NeuN*^POS^* ChAT*^NEG^*and on-target NeuN*^POS^* ChAT*^POS^* spinal motor neurons as described in Figure 5. Individual cells are plotted for saline, ΔCRE controls, mouse CRE98, and CAG (n=3 animals, data reproduced from Figure 3) and test constructs (n=2 animals for 3mCORE, otherwise n=3 animals). Data are separated by animal. Mean intensity across all cells per spinal cord section denoted by white circle, distribution of all cells by section denoted by box-and-whisker plot (median denoted by white line, first and fourth quartiles by vertical line, outliers by black points). **(c)** Representative IHC images of T1-L4 DRG simultaneously dissected from previously described animals (Figures 3 and 5) 14 days after P0 ICV injection of candidate CRE reporter AAVs (DAPI, cyan; GFP, yellow). Images for mCRE98 and CAG reproduced from Figure 3e. Images acquired at pBG promoter-optimized acquisition parameters. Scale bar 100 μm. Percentage of GFP*^POS^***(d)** and mean GFP intensity **(e)** in off-target DRG neurons (visually identified by nuclear DAPI morphology and size) for saline and ΔCRE controls, CRE98, synthetic and human CRE98 constructs, and CAG (n=2 animals for 3mCORE, otherwise n=3 animals). Mean percentage across all DRG sections per animal denoted by each point, mean across all animals per condition denoted by bar plot, standard deviation across all animals denoted by line. Images acquired and quantified at pBG promoter-optimized acquisition parameters. Data for saline, ΔCRE, mCRE98, and CAG constructs reproduced from Figure 3c).

